# Dietary magnesium supplement enhances memory through Hippo signaling

**DOI:** 10.64898/2026.10.02.756214

**Authors:** Yanying Wu, Christoph D. Treiber, Scott Waddell

## Abstract

Dietary magnesium (Mg^2+^) supplement enhances memory in mammals and *Drosophila*, but whether and how Mg^2+^ changes molecular states of neurons, remains unknown. Here we used single-cell RNA sequencing to map cell-type specific transcriptional responses to memory-enhancing dietary Mg^2+^. These analyses revealed changes in Hippo-pathway gene expression in memory-relevant αβ Kenyon Cells (KCs) of the mushroom bodies. Targeted knock-down of *Merlin*, *hippo* (*hpo*) and *warts* demonstrated requirement in αβ KCs for Mg^2+^-enhanced, but not baseline memory. Further study of Hippo signaling identified Yorkie (*yki*) transcriptional coactivator and DNA-binding partner Scalloped (*sd*) to be required for baseline and Mg^2+^-enhanced memory. Moreover, releasing Yki::Sd of inhibition from Tondu-domain containing growth inhibitor (*Tgi*) specifically in αβ KCs enhanced memory, independent of diet. Analyses of Yki::Sd target genes identified several required in αβ KCs for baseline memory, including *crb* and *Glut1* which are also Mg^2+^-induced in a Hippo-dependent manner in αβ KCs. These genes suggest that aiding the metabolic demands of long-term memory and facilitating structural neuronal plasticity are key mediators of Mg^2+^-induced memory enhancement. We propose pharmacological enhancement of Sd-dependent transcription, and activity of particular Sd-target genes may have therapeutic potential for memory disorders.

## Introduction

Health food store shelves are stacked with nootropics, a diverse group of supplements that claim to improve learning and memory. Few of these ‘smart drugs’ or ‘cognitive enhancers’ – often plant extracts, vitamins and minerals – have been experimentally validated, making their efficacy questionable. An interesting exception amongst nootropics is dietary magnesium (Mg^2+^) supplementation, which has been reported to enhance neuronal plasticity and memory performance of young and aged rodents, measured in a variety of behavioral tasks (*1–4*). In addition, dietary elevation of brain Mg^2+^ enhanced retention of the fear extinction memories (*5*). The beneficial effects indicate dietary Mg^2+^ may have therapeutic value for patients with a variety of memory-related problems (*6*).

The memory-enhancing property of Mg^2+^ in rodents was largely attributed to increased synaptic density in the dentate gyrus and CA1 subregions of the hippocampus, and activity of N-methyl-D-aspartate type glutamate receptors (NMDAR) (*2*, *7*). However, Mg^2+^ is a co-factor for hundreds of enzymes (*8*, *9*), making it likely that other molecular sites of action could facilitate memory.

Dietary Mg^2+^ supplement also enhances long-term memory (LTM) in *Drosophila* (*10*), suggesting evolutionarily conserved signaling pathways may underlie Mg^2+^-dependent memory enhancement. Importantly, *Drosophila* permits genetic approaches to decipher mechanisms underpinning memory enhancement. In our initial study we found that memory-enhancing levels of dietary Mg^2+^ elevated intracellular Mg^2+^ in memory-relevant αβ Kenyon Cells (KCs) of the mushroom bodies (*10*). Moreover, memory enhancement in the fly is independent of NMDARs and instead depends on αβ KC expression of a CNNM-type magnesium efflux transporter encoded by the *unextended* (*uex*) gene (*10*). Together these findings indicated that αβ KC-specific adaptation to elevated Mg^2+^ may underly memory enhancement. Here we identify changes in Hippo signaling as one of these adaptations and for cell-type specific manipulations of Hippo-dependent gene expression to be memory enhancing independent of diet.

## Results

### Single-cell transcriptomic analyses of dietary Mg^2+^ supplementation

Although it is broadly appreciated that diet alters body-wide gene expression (*11*), we previously showed memory enhancing effects of dietary Mg^2+^ to specifically involve Mg^2+^ transport in αβ KCs in the fly’s mushroom bodies (*10*). We therefore used 10x Genomics Chromium single-cell RNA sequencing (scRNA-seq) to investigate how the fly brain, and particularly αβ KCs, adapt to memory-enhancing levels of dietary Mg^2+^. In earlier work we established 80 mM Mg^2+^ supplement to produce optimal LTM enhancement (*10*). We therefore *ad libitum* fed flies for 4 days with food containing either 1 mM MgCl_2_ (low [Mg^2+^]) or supplemented with 80 mM MgCl_2_ (high [Mg^2+^]) (Figure 1A). Each group contained four biological replicates comprising independent groups of flies. We prepared scRNA-seq libraries from central brains from 10 male and 10 female flies from each of 8 independent groups; 4 low [Mg^2+^] and 4 high [Mg^2+^].

**Figure 1.**
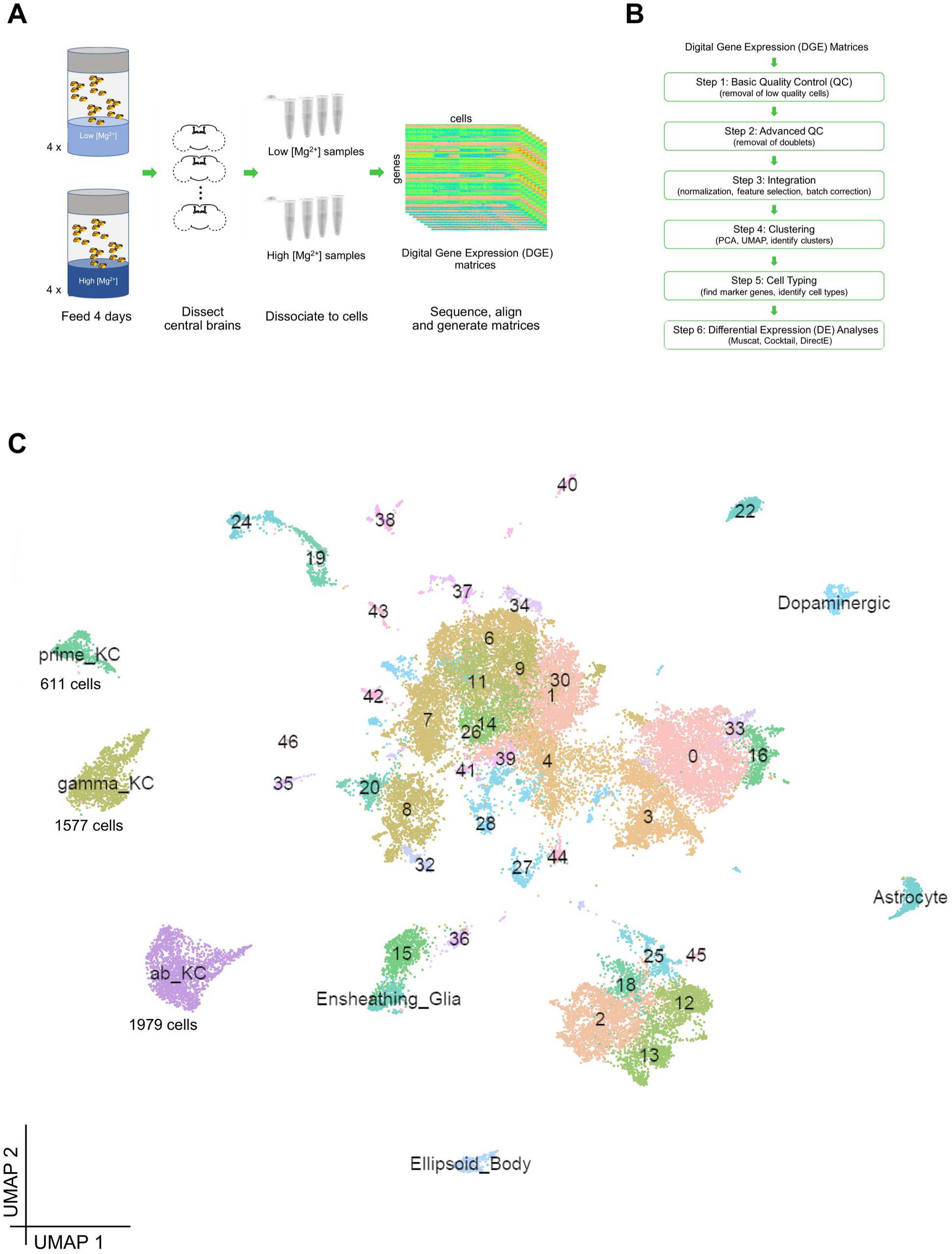
Clustering and identification of major cell types. (**A**) Protocol for conditional scSeq experiment. 8 groups of flies were fed either high (80 mM) or low (1 mM) [Mg^2+^] food for 4 days before their central brains were dissected and dissociated to cell suspensions. 4 high [Mg^2+^] and 4 low [Mg^2+^] samples were sequenced using 10x Genomics technology. Raw sequences were aligned to the *Drosophila* genome (R6.25) to produce 8 digital gene expression (DGE) matrices. (**B**) Workflow for data processing. DEG matrices were subjected to quality control (QC) to remove low-quality cells and doublets (step 1-2), then integrated, clustered and annotated for cell types (step 3-5). Finally, differential expression (DE) analyses compared transcriptional profiles between high and low [Mg^2+^] samples (step 6). (**C**) A uniform manifold approximation and projection (UMAP) visualization of 38,414 single cells, clustered into 47 groups. 7 major cell types were annotated using known marker genes: αβ KCs (ab_KC), α′β′ KCs (prime_KC), γ KCs (gamma_KC), Dopaminergic neurons (Dopaminergic), Astrocytes (Astrocyte), Ensheathing Glia (Ensheathing_Glia) and Ellipsoid Body ring neurons (Ellipsoid_Body). Numbers of cells in each KC cluster are shown.

Sequencing results were subsequently aligned to the fly genome to generate 8 digital gene expression (DGE) matrices, which were then analyzed using the workflow summarized in Figure 1B (see materials and methods for details). Briefly, after filtering out low-quality cells and putative cell doublets we retrieved 38,414 high-quality cells for further analyses (Figure 1B, Step 1 and 2; Supplementary Figures 1 and 2). SCTransform and the canonical correlation anchor-based method in Seurat v3 (*12*, *13*) were used to normalize and integrate data from different samples (Figure 1B, Step 3). We subsequently identified 47 cell clusters using a modularity optimization algorithm based on a shared nearest neighbor measurement (*14*) (Figure 1B, Step 4) and visualized these clusters with Uniform Manifold Approximation and Projection (UMAP) (Figure 1C) (*15*). Importantly, each experimental sample contributed a comparable proportion of cells to every cluster (Supplementary Figure 3, A and C). Among the 47 clusters, we easily identified the 3 major KC types of the mushroom bodies (*16*, *17*): αβ (ab_KC, 1979 cells), α′β′ (prime_KC, 611 cells) and γ KCs (gamma_KC, 1577 cells) and dopaminergic neurons, Ellipsoid Body (ring neurons), Astrocytes and Ensheathing Glia (Figure 1C), using expression of known marker genes.

**Figure 2.**
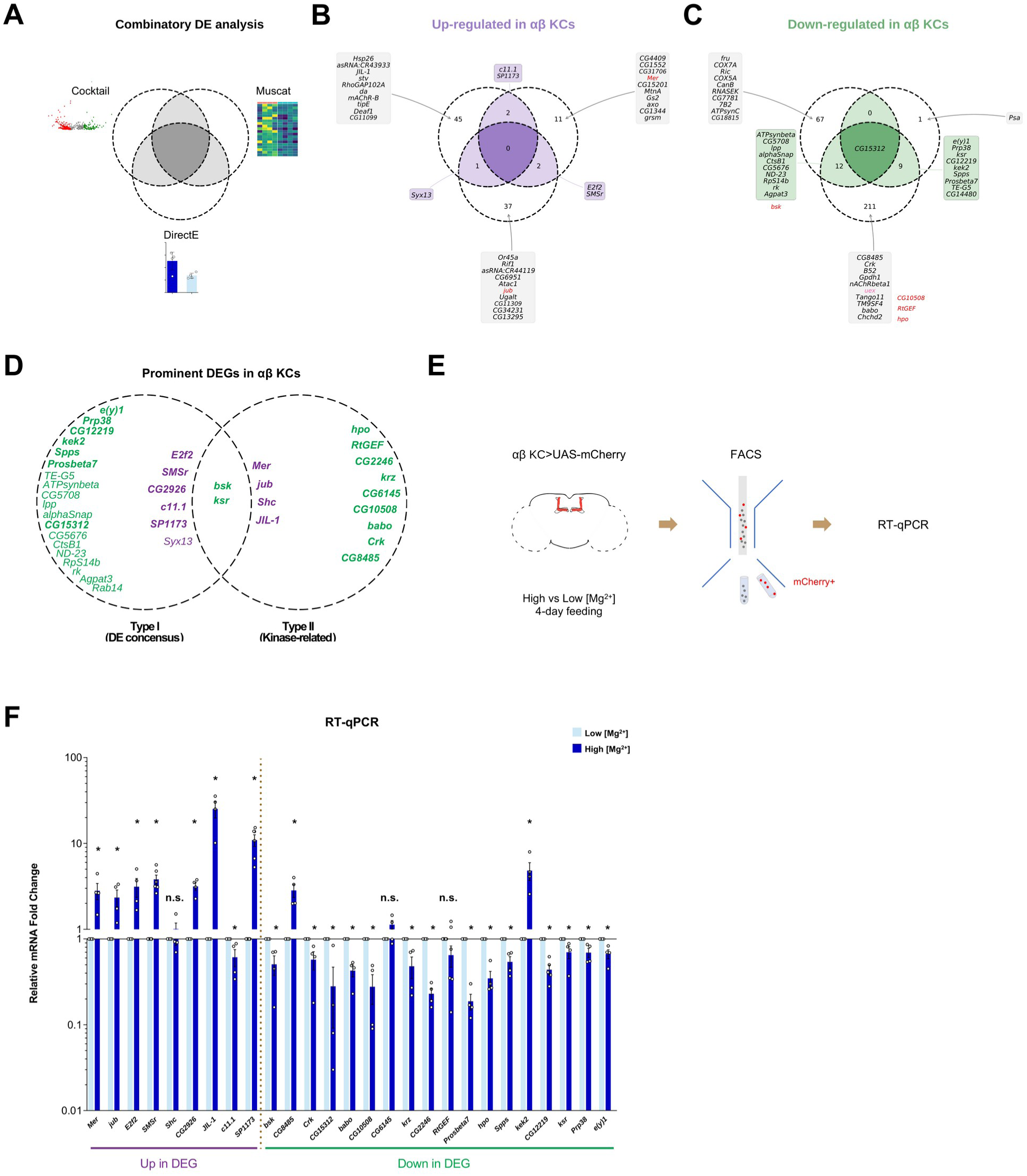
High [Mg^2+^] diet changes gene expression in αβ Kenyon Cells. (**A**-**C**) Venn plots of integrated DE results from the 3 methods showing (B) up-regulated and (C) down-regulated DEGs in αβ KCs. Each region of the Venn plot shows corresponding numbers of DEGs, except for *CG15312* in center of (C), and list of genes. If DEG > 10, only the top 10 are displayed. Circles are arranged as in (A): Cocktail (upper left), Muscat (upper right) and DirectE (bottom). Hippo pathway-related DEGs (*Mer*, *jub*, *bsk*, *CG10508*, *RtGEF* and *hpo*) are shown in red; those outside the displayed top 10 (*bsk*, *CG10508*, *RtGEF* and *hpo*) are noted beside the relevant region. *uex* is shown in pink. (D) Venn plot of filtered DEGs in αβ KCs. 27 Type I DEGs, genes identified in at least 2 of 3 DE methods, and 15 ‘kinase-related’ Type II DEGs are shown. Two genes, *bsk* and *ksr*, are common. Up-regulated DEGs (purple) and down-regulated DEGs (green) are marked, with those subjected to RT-qPCR validation in bold. (E) Protocol of RT-qPCR validation. Flies expressing mCherry in αβ KCs were fed high or low [Mg^2+^] diet for 4 days. Central brains were dissected, dissociated, and αβ KCs (mCherry positive cells) were purified from cell suspensions using FACS. mRNAs extracted from sorted cells were reverse transcribed and amplified for qPCR measurement. (F) Relative mRNA fold changes of DEGs in αβ KCs from flies fed high [Mg^2+^] versus low [Mg^2+^] diet. Each data point represents average of 3 technical replicates from one biological replicate (n = 1). \**P* < 0.05, *t* test, *n* = 4-6.

### Mg^2+^ supplemented diet produces cell-type specific changes of gene expression

Since dietary Mg^2+^ should be available to every cell, we first asked how broadly it alters gene expression by applying the Muscat DE analysis to the seven major cell types listed above (Supplementary Figure 4, A and B). Each cell type responded to high [Mg^2+^] diet with a unique set of DEGs (Supplementary Figure 4C and Supplementary Data 1). Perhaps surprisingly, Astrocytes, Ensheathing Glia and Ellipsoid Body neurons were skewed towards down-regulation, with more DEGs at log_2_FC < −1 and fewer at log_2_FC > 1 than the other cell types.

Overlap between cell types was minimal, with the most notable shared response being upregulation in αβ, α′β′ and γ KCs of *CG1552*, encoding an arthropod-specific SVC-family protein implicated in environmental stress (*18*) (Supplementary Figure 4D). The most strongly regulated genes of each cell type are shown in Supplementary Figure 4F. Dietary Mg^2+^ supplement therefore alters gene expression throughout the brain but in a cell-type-specific manner.

### Differential Mg^2+^-dependent expression in Kenyon Cells

We next performed a more detailed analysis of DEGs between KC types, given their distinct roles in memory processing (*19–23*). Here we took a consensus approach of combining three DE methods, Cocktail, Muscat and DirectE (*24–27*), to generate a list of candidate DEGs, for validation and functional testing using the unique strength of *Drosophila* genetics (Figure 1B, Step 6 and Figure 2A and Supplementary Data 2; see materials and methods for details). Cocktail (Supplementary Figure 5A and Supplementary Data 3), Muscat (Supplementary Figure 5, B and C) and DirectE (Supplementary Figure 5, D and E) reported largely different lists of αβ KC DEGs. Together they returned 98 up-regulated and 301 down-regulated DEGs (FDR adjusted *P* < 0.05) (Figure 2, B and C; Supplementary Data 2). Only 5 up-regulated (*c11.1*, *sp1173*, *Syx13*, *E2f2* and *SMSr*) and 22 down-regulated genes (Figure 2, B and C) were identified by two or three of the methods. As comparison, DE analyses identified 204 up-regulated and 93 down-regulated DEGs in α′β′ KCs (Supplementary Figure 6), whereas γ KCs exhibited 134 up-regulated and 323 down-regulated DEGs (Supplementary Figure 7). All 3 DE methods, and the combined lists, show that each KC type responded differently to dietary [Mg^2+^] supplement (Supplementary Data 2), with overlap between the DEGs in each KC subtype being rare (Supplementary Figure 8). Therefore, the distinct KC types also respond in unique ways to dietary Mg^2+^ supplement, which could differentially contribute to memory enhancement.

### Selection and validation of differentially expressed αβ Kenyon Cell genes

We focused functional analyses on αβ KC expressed genes, given the established importance of this cell type for normal and Mg^2+^-enhanced LTM (*10*, *19*, *20*, *28*, *29*). Our combined DE analyses of αβ KCs identified 399 candidate Mg^2+^-regulated DEGs, which we streamlined for molecular validation and functional testing by defining two categories.

’Type I’ genes were those detected as being differentially expressed in αβ KCs by at least 2 of the 3 DE methods. This filtering returned 27 genes; 6 up-regulated (marked purple in Figure 2D, left circle) and 21 down-regulated (marked green in Figure 2D, left circle).

’Type II’ DEGs were defined by their functional category identity in FlyBase (GENE_SNAPSHOT, GO_BP, GO_CC, GO_MF) (*30*, *31*) as ‘kinase’, given that kinases represent a prominent class of Mg^2+^-dependent enzymes and play pivotal roles in cell-signaling cascades (*32*, *33*). Applying the ‘kinase’ filter yielded 22 ‘kinase-related’ DEGs (Supplementary Data 4), which showed enrichment for components of the mitogen-activated protein kinase (MAPK) (six: *bsk*, *Crk*, *hpo*, *krz*, *ksr* and *shc*) and Hippo (six: *bsk*, *CG10508*, *jub*, *hpo*, *Mer* and *RtGEF*) signaling pathways. Type II filtering also returned 5 other kinases that are not evidently linked to these pathways. Therefore, in total we defined 15 ‘Type II’ DEGs (2 genes, *bsk* and *hpo*, feature in both the MAPK and Hippo pathways). Four of these 15 ‘Type II’ genes were up-regulated in the high-Mg^2+^ feeding group (Figure 2D, right circle), whereas eleven were down-regulated (Figure 2D, right circle).

Combining ‘Type I’ and ‘Type II’ lists produced 40 candidate DEGs (Figure 2D, Supplementary Data 5), from which 27 were selected for validation using RT-qPCR of mRNAs extracted from FAC-sorted mCherry-labeled αβ KCs isolated from flies fed low or high [Mg^2+^] diet (Figure 2E, see materials and methods for details). The 27 genes comprised the top 5 up-regulated and top 6 down-regulated Type I genes together with all Type II genes (Figure 2D and Supplementary Data 5), and also *CG15312* because it was the only gene detected by all three DE methods (Figure 2C). RT-qPCR confirmed differential expression of 21 of these 27 DEGs in FAC-sorted αβ KCs, compared to their relative expression in analyses of scRNA-seq data (Figure 2F).

Differential expression of *shc*, *CG6145* and *RtGEF* was not validated, while expression of *c11.1*, *CG8485* and *kek2* changed in the opposite direction to that reported by scRNA-seq data (Figure 2F). We therefore removed these 6 genes from further consideration.

We also noted that DirectE identified *uex*, the founding gene of our Mg^2+^ and memory studies (*10*), to be down-regulated in αβ KCs (Figure 2C and Supplementary Data 6), which we confirmed using our previously constructed *uex::HA* gene tag fly strain (*10*) (Supplementary Figure 9). *uex* encodes the fly CNNM-type Mg^2+^ efflux transporter and is required for both normal and Mg^2+^-enhanced memory (*10*). We were previously unable to reconcile finding that intracellular [Mg^2+^] is elevated in αβ KCs of flies fed memory-enhancing [Mg^2+^] when UEX mediates Mg^2+^ efflux. However, *uex* down-regulation in these cells should reduce Mg^2+^ efflux and thereby increase intracellular [Mg^2+^] in flies fed memory-enhancing diet.

### αβ KC expression of *Merlin* and *Hippo* is required for memory enhancement

We next tested the functional importance in Mg^2+^-enhanced memory of each of the validated αβ KC regulated genes by knocking down their expression using transgenic RNA interference (RNAi). Flies expressing c739-GAL4 driven UAS-RNAi constructs targeting either *Mer*, *CG2926*, *JIL-1*, *bsk*, *CG2246* (phosphoribosyl pyrophosphate synthetase associated protein, PRPSAP), *hpo*, *Spps*, *CG12219*, *ksr*, and *Prp38* in the αβ KCs (Supplementary Figure 10) exhibited similar memory performance when fed with either low or high [Mg^2+^]. Loss of diet-dependent memory enhancement was replicated when RNAi for these genes was driven using the more αβ KC-restricted MB008B-GAL4 driver (*16*, *34*) (Figure 3A). Although all manipulations abolished Mg^2+^-dependent memory enhancement, we noted that some appeared to enhance memory compared to genetic controls (*CG2246*, *Spps*, *CG12219* and *ksr*), whereas others impaired it (*CG2926*, *JIL-1*, *bsk* and *Prp38*), producing similarly high or low LTM performance in flies fed either diet.

**Figure 3.**
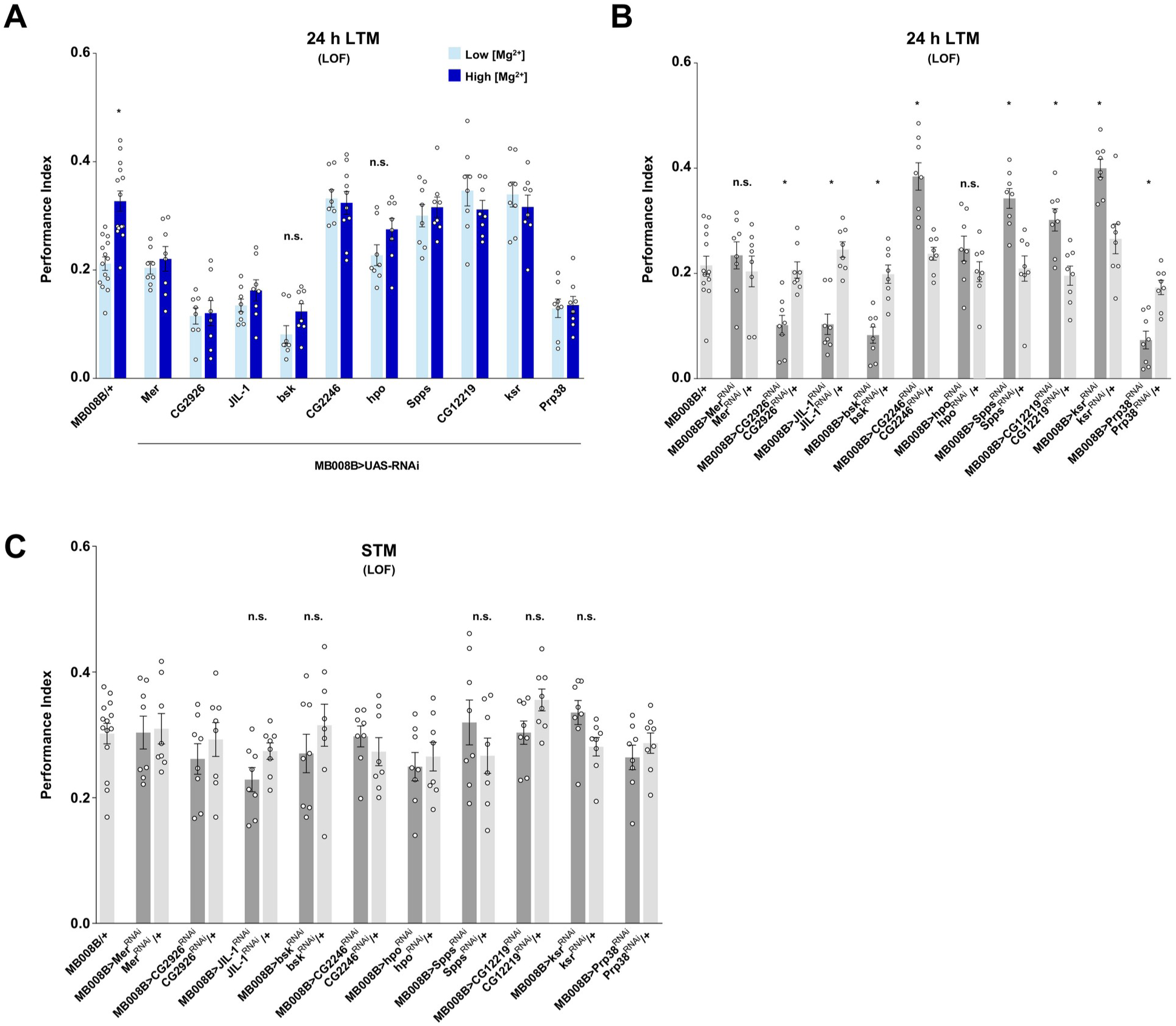
Prominent DEGs are differentially required in αβ KCs for memory enhancement. (**A**) The memory enhancing effect of high [Mg^2+^] diet is lost in flies expressing RNAi targeting several prominent DEGs in αβ KCs (driven by MB008B-GAL4). Their 24 h appetitive LTM was either similarly low or high in flies fed either diet (\**P* < 0.05, *t* test, *n* = 8-14). (**B**) LTM of flies fed low [Mg^2+^] diet was significantly impaired in flies harboring αβ KC knockdown of *CG2926*, *JIL-1*, *bsk* or *Prp38*, while it was significantly enhanced in those with *CG2246*, *Spps*, *CG12219* or *ksr* knockdown, as compared to controls. No significant difference of LTM was observed when knocking down *Mer* or *hpo* (\**P* < 0.05, ANOVA, *n* = 8-14). (**C**) Short-term memory (STM) was not affected by knockdown of these genes (ANOVA, *n* = 8-14).

To better understand the mnemonic effects of the gene manipulations, we tested whether they altered LTM in flies fed low [Mg^2+^] diet. We again knocked down gene expression in αβ KCs (MB008B-GAL4) and assayed appetitive memory. Short-term memory (STM) measured immediately after training was not significantly affected by any of the manipulations (Figure 3C). However, gene-specific effects were observed for LTM (Figure 3B). Whereas flies expressing RNAi for *CG2926*, *JIL-1*, *bsk* or *Prp38* exhibited impaired LTM, those harboring αβ KC knockdown of *CG2246*, *Spps*, *CG12219* or *ksr* showed enhanced LTM, resembling that observed with high [Mg^2+^] diet (Figure 3B). Surprisingly, flies expressing *Mer* or *hpo* RNAi in αβ KCs exhibited LTM that was no different to that of their relevant controls (Figure 3B). We therefore concluded that the Hippo pathway components Mer and Hpo were unique in being specifically required for Mg^2+^-dependent memory enhancement (Table 1).

**Table 1.** Test result summary for the 10 DEGs impacting memory enhancement. First column shows the gene names. Second column shows the DE analysis results from the scRNA-seq data. An up-arrow indicates that the expression of the gene was up-regulated in the high [Mg^2+^] sample compared to the low [Mg^2+^] sample, and a down-arrow means the gene was down-regulated. Third column shows the RT-qPCR test results, arrows mean the similar up or down-regulation of the gene as column 2. Fourth, fifth and sixth columns show the behavior results extracted from Figure 3A, 3B and 3C, respectively. An up-arrow indicates a significantly higher performance, a down-arrow denotes a significantly lower performance and an ‘n.s.’ means no significant difference in behavior performance comparing the relevant groups. Last column, a star indicates that the corresponding gene shows an enhancement-specific role. Rows are grouped by phenotype class: enhancement-specific genes, genes whose knockdown impairs baseline LTM, and genes whose knockdown enhances baseline LTM.

| Gene | scRNA-seq<br>DE<br>$\alpha\beta$ KCs<br>High vs Low<br>[Mg<sup>2+</sup>] | RT-qPCR<br>$\alpha\beta$ KCs<br>High vs Low<br>[Mg<sup>2+</sup>] | 24-hour<br>memory<br>RNAi, High vs<br>Low [Mg<sup>2+</sup>]<br>(wild type ↑) | 24-hour<br>memory<br>RNAi vs<br>control | Immediate<br>memory<br>RNAi vs<br>control | Enhancement<br>specific |
| --- | --- | --- | --- | --- | --- | --- |
| **Enhancement-specific** |  |  |  |  |  |  |
| *Mer* | ↑ | ↑ | n.s. | n.s. | n.s. | * |
| *hpo* | ↓ | ↓ | n.s. | n.s. | n.s. | * |
| **Knockdown impairs baseline LTM** |  |  |  |  |  |  |
| *CG2926* | ↑ | ↑ | n.s. | ↓ | n.s. |  |
| *JIL-1* | ↑ | ↑ | n.s. | ↓ | n.s. |  |
| *bsk* | ↓ | ↓ | n.s. | ↓ | n.s. |  |
| *Prp38* | ↓ | ↓ | n.s. | ↓ | n.s. |  |
| **Knockdown enhances baseline LTM** |  |  |  |  |  |  |
| *CG2246* | ↓ | ↓ | n.s. | ↑ | n.s. |  |
| *Spps* | ↓ | ↓ | n.s. | ↑ | n.s. |  |
| *CG12219* | ↓ | ↓ | n.s. | ↑ | n.s. |  |
| *ksr* | ↓ | ↓ | n.s. | ↑ | n.s. |  |

### Hippo-regulated transcription is critical for memory enhancement

The Mg^2+^ diet-selective memory phenotypes of *Mer* and *hpo* loss of function led us to focus on a potential role for Hippo signaling in Mg^2+^-dependent memory enhancement. The conserved Hippo pathway involves a core kinase cascade of Hippo (*hpo*, a Ser/Thr protein kinase; mammal Sterile 20-like kinases 1 and 2, MST1/2) and its regulator Salvador (*sav*; mammal SAV1 or WW45, WWP4), which together phosphorylate and activate Warts (*wts*, also a Ser/Thr protein kinase; mammal LATS1/2). Wts in turn, along with its regulator Mob as tumor suppressor (*mats*; mammal MOB1A/B), phosphorylates and inactivates Yorkie (*yki*; mammal YAP, Yes-associated protein and TAZ, transcriptional co-activator with PDZ-binding motif), an oncoprotein and transcriptional co-activator, which associates with the DNA binding transcription factor Scalloped (*sd*; mammal TEAD1-4). Tondu-domain-containing Growth Inhibitor (*Tgi*, mammal Vestigial-like 4, VGLL4) competes with Yki for binding to Sd and thereby inhibits Yki::Sd-dependent gene expression (*35*, *36*). Merlin (*Mer*; mammal neurofibromatosis 2, NF2) and Expanded (*ex*; mammal expanded, EX) are upstream activators of Hippo (*35*, *37–40*).

We used UAS-RNAi to knockdown expression of these core Hippo pathway genes specifically in αβ KCs and tested their roles in memory enhancement (Figure 4A). In addition, since Hippo signaling plays critical roles in development (*39*, *41*, *42*), we restricted gene knockdown to adulthood using GAL80^ts^ (*43*). At permissive 18°C, GAL80^ts^ binds to GAL4 and inhibits its transcriptional activator function. This suppression is released, and the UAS-RNAi transgene is expressed, at restrictive 30°C. Of the nine Hippo pathway genes tested, knockdown of six—*Mer*, *hpo*, *wts*, *Tgi*, *yki*, and *sd*—abolished the memory enhancing effect of high [Mg^2+^] diet, while knockdown of *ex*, *sav*, and *mats* had no effect (Figure 4B). Mg^2+^-dependent memory enhancement remained intact if RNAi transgenes were not induced (Supplementary Figure 11A), confirming that loss of enhancement resulted from acute gene knockdown in adult αβ KCs. Moreover, no defects were observed when flies were tested for task-relevant sensory acuity (i.e. odor and sugar sensitivity; Supplementary Data 7).

**Figure 4.**
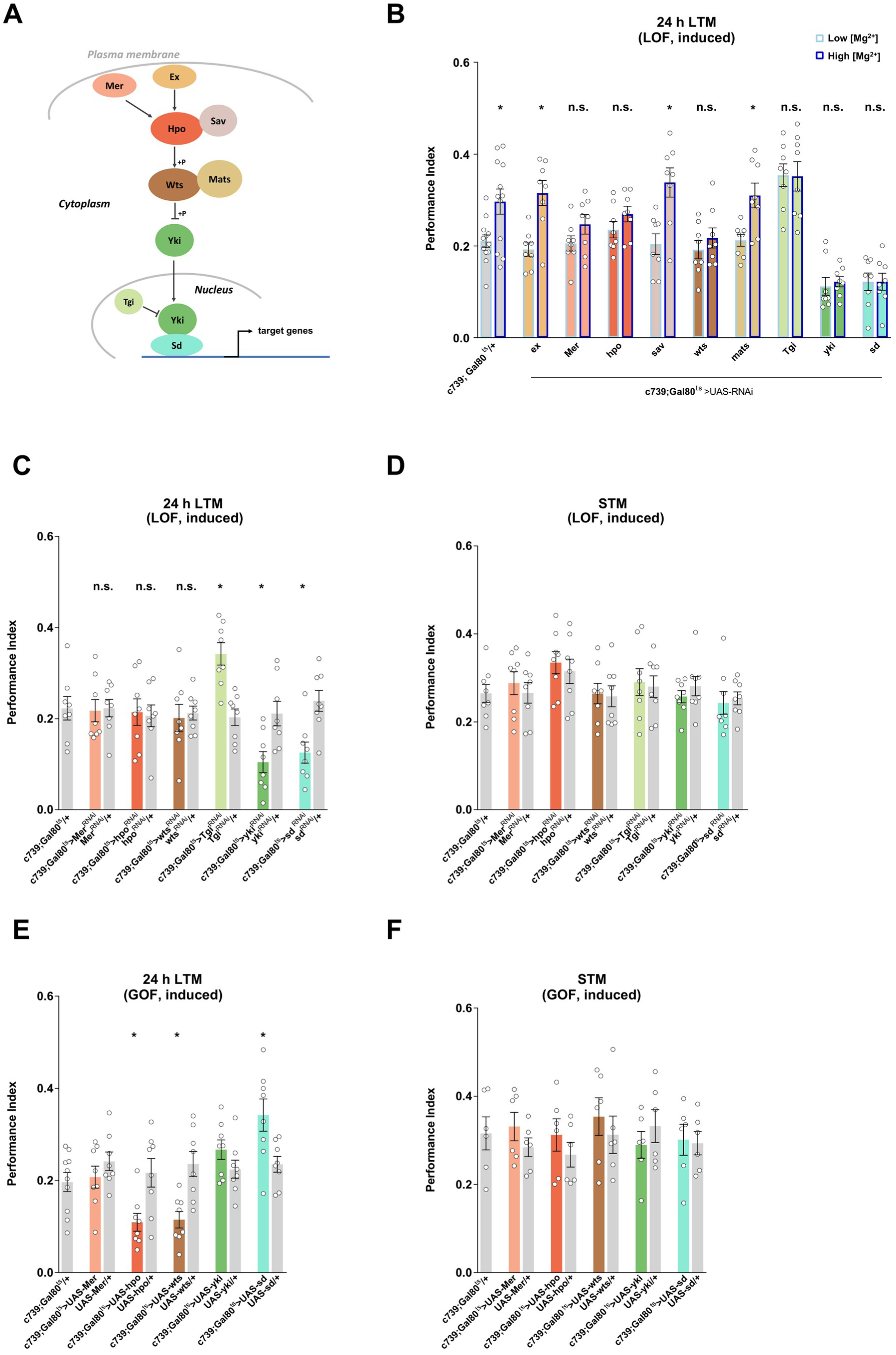
Hippo pathway genes are central to memory enhancement. (**A**) Schematic of core Hippo signaling. (**B**) Adult restricted expression in αβ KCs of RNAi targeting *Mer*, *hpo*, *wts*, *Tgi*, *yki* or *sd* abolished [Mg^2+^] diet-dependent enhancement of LTM, while flies with knockdown of *ex*, *sav*, or *mats* retained diet-dependent enhancement (\**P* < 0.05, *t* test, *n* = 8-10). (**C**) LTM was significantly impaired when *yki* or *sd* RNAi knockdown was restricted to adult αβ KCs of flies fed low [Mg^2+^] diet, but not when *Mer*, *hpo*, or *wts* were targeted. In contrast, knockdown of *Tgi* in adult αβ KCs significantly enhanced LTM (\**P* < 0.05, ANOVA, *n* = 8). (**D**) STM was not affected in flies with adult αβ KC-restricted knockdown of *Mer*, *hpo*, *wts*, *Tgi*, *yki* or *sd* (ANOVA, *n* = 8). (**E**) LTM was significantly impaired in flies fed low [Mg^2+^] diet when *hpo* or *wts* were overexpressed in αβ KCs of adult flies, while it was significantly enhanced by *sd* overexpression (\**P* < 0.05, *t* test, *n* = 8-10). (**F**) STM was not affected by adult αβ KC overexpression of *Mer*, *hpo*, *wts*, *yki* or *sd* (ANOVA, *n* = 6).

We also tested roles in memory for these Hippo pathway genes in flies fed low [Mg^2+^] diet. Knockdown of *Mer*, *hpo*, or *wts* in adult αβ KCs had no effect (Figure 4, C and D), consistent with our initial *Mer* and *hpo* experiments (Figure 3, A and B), and confirming these genes to be uniquely required for Mg^2+^-dependent memory enhancement. In contrast, adult-restricted *sd* or *yki* knockdown in αβ KCs also significantly impaired LTM in flies fed low [Mg^2+^] diet (Figure 4, C and D). Again, no LTM effects were evident if RNAi transgenes were not induced (Supplementary Figure 11B). Importantly, STM was not affected in *sd* and *yki* knockdown flies (Figure 4D), revealing a novel and LTM specific requirement for Yki and Sd.

Since Tgi is a competitive inhibitor of Sd::Yki association, we hypothesized that releasing its influence might enhance LTM independently of Mg^2+^ diet. Indeed, adult-restricted knockdown in αβ KCs of *Tgi* significantly enhanced LTM (Figure 4C). In contrast, adult αβ KC restricted knockdown of nervous fingers 1 (*nerfin-1*, mammal INSM transcriptional repressor 1), another competitive inhibitor of Sd::Yki association that binds to a different region of Sd than Tgi (*44*, *45*), did not enhance or impair LTM (Supplementary Figure 11F). To corroborate that Tgi-directed gain of function resulted from Yki::Sd dependent transcription, we tested LTM in flies overexpressing UAS-driven transgenes for *Mer*, *hpo*, *wts*, *sd*, and *yki*. Whereas overexpression in adult αβ KCs of *hpo* or *wts* significantly and specifically impaired LTM, overexpression of *sd* enhanced LTM and *yki* had no obvious effect (Figure 4, E and F), and LTM was unaffected if the transgenes were not induced (Supplementary Figure 11E). VGLL4 (Tgi) is a known tumor suppressor, so no pharmaceuticals have been developed to inhibit its function. We therefore tested whether feeding IK-930, a potent, orally active inhibitor of TEAD1 (human ortholog of Sd) (*46*), affected LTM. Flies fed for 4 days with food containing 10, 20 or 40 μM IK-930 exhibited a dose-dependent impairment of LTM (Supplementary Figure 11D), which provides further support for an acute role of Sd in LTM. In contrast, feeding flies for 4 days with 10, 20 or 40 μM XMU-MP-1, an inhibitor of the Hpo orthologs MST1/2 (47, 48), did not affect LTM (Supplementary Figure 11C), consistent with *hpo* knockdown leaving baseline LTM intact (Figure 4C).

In summary, our manipulations of core Hippo pathway components revealed roles for *Mer*, *hpo*, and *wts* in Mg^2+^-dependent memory enhancement, and for *Tgi*, *yki* and *sd* in both Mg^2+^-dependent memory enhancement and diet-independent LTM.

### Sd/TEAD target genes *crb* and *Glut1* are effectors of memory enhancement

Sd acts as an effector of the Hippo pathway by driving transcription of target genes. We therefore further investigated how Sd affects LTM by testing roles for the known Sd target genes *expanded* (*ex*), *Death-associated inhibitor of apoptosis 1* (*Diap1*), *Cyclin E* (*CycE*), *Myc* (*Myc*), *kibra* (*kibra*), *four-jointed* (*fj*), *crumbs* (*crb*), *division abnormally delayed* (*dally*), *wingless* (*wg*) (*37*, *38*, *41*, *42*) in diet-independent and Mg^2+^-enhanced LTM. We also included *Glut1*, because its human ortholog, *GLUT3*, was identified as a YAP/TAZ/TEAD regulated gene (*49*). RNAi-mediated knockdown in adult αβ KCs of *kibra*, *crb*, *wg*, or *Glut1* significantly impaired LTM in flies fed low [Mg^2+^] diet, while *ex*, *Diap1*, *CycE*, *Myc*, *fj*, and *dally* RNAi had no effect (Figure 5, A and B, and Supplementary Figure 12A; sensory acuity, Supplementary Data 8). No LTM defects were evident when these RNAi transgenes were not induced (Supplementary Figure 12B). Therefore, these data indicate potential roles for a subset of Sd regulated genes, *kibra*, *crb*, *wg*, or *Glut1*, in LTM. Mushroom body knockdown of *wg* was previously shown to impair aversive LTM (*50*).

**Figure 5.**
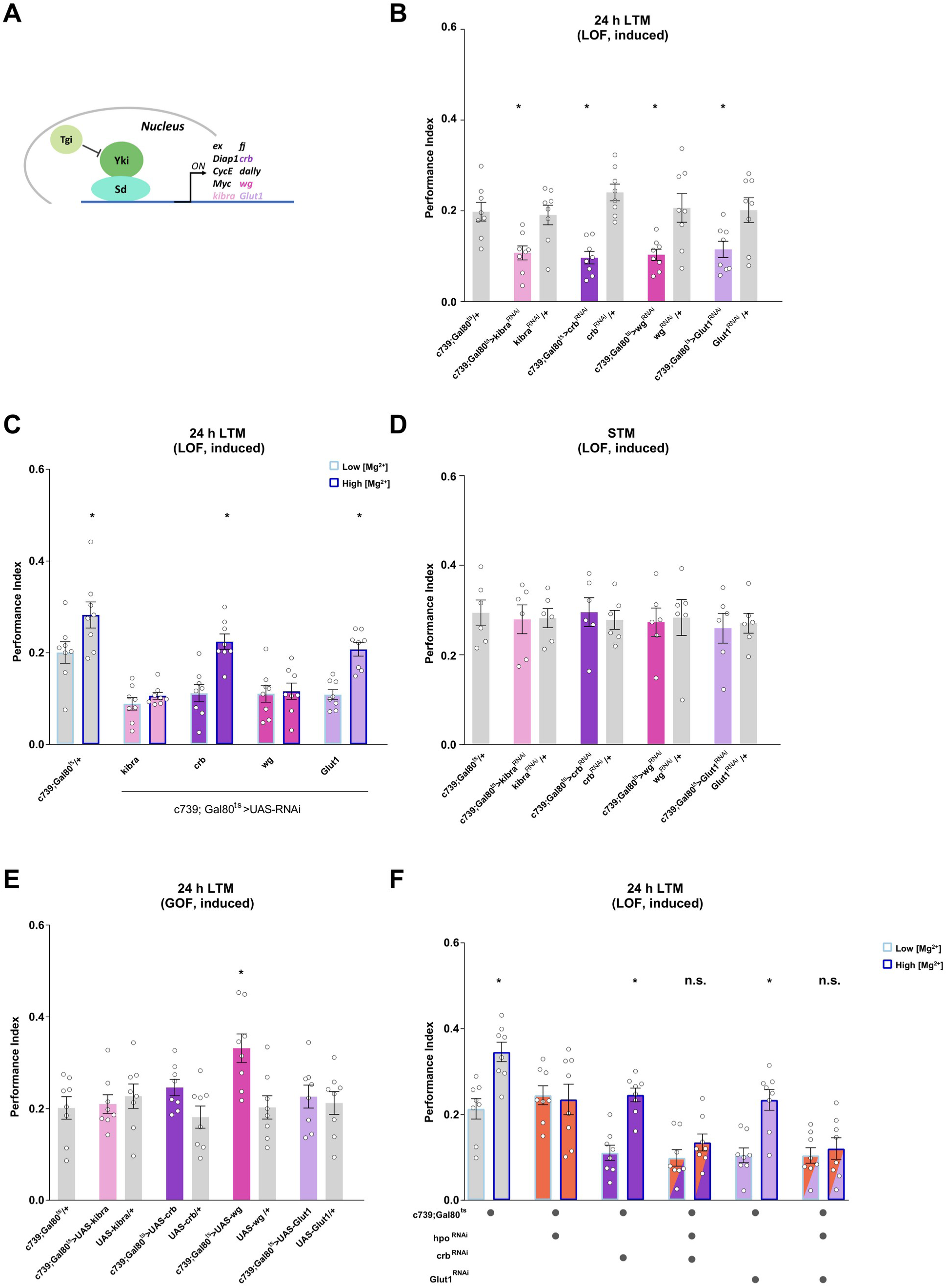
Hippo pathway targets *crb* and *Glut1* are effectors of memory enhancement. (**A**) Illustration of Yki::Sd activation of target genes. Tgi inhibits Yki binding to Sd. (**B**) LTM was impaired when *kibra*, *crb*, *wg* or *Glut1* RNAi knockdown was restricted to adult αβ KCs of flies fed low [Mg^2+^] diet (\**P* < 0.05, ANOVA, *n* = 8). (**C**) Defective LTM of flies harboring *crb* or *Glut1* knockdown in adult αβ KCs was rescued by high [Mg^2+^] diet, while defective performance of *kibra* and *wg* knockdown flies remained (\**P* < 0.05, *t* test, *n* = 8). (**D**) STM was unaffected by RNAi manipulation of Yki::Sd target genes (ANOVA, *n* = 6). (**E**) LTM was enhanced in flies overexpressing *wg* in αβ KCs of adult flies, while it was unaffected by *kibra*, *crb* or *Glut1* overexpression (\**P* < 0.05, *t* test, *n* = 8). (**F**) High [Mg^2+^] diet could not rescue LTM of flies harboring adult αβ KC knockdown of *crb* or *Glut1* when *hpo-RNAi* was co-expressed (\**P* < 0.05, *t* test, *n* = 8).

Finding that *hpo* was downregulated by high [Mg^2+^] diet (Figure 2, D and F) led us to reason that a resulting reduction of Wts and Yki phosphorylation should allow Yki to promote Sd-dependent gene expression (Figure 4A). We therefore tested whether Mg^2+^-dependent upregulation of relevant Sd target genes might counteract LTM defects produced by their RNAi-mediated knockdown. We compared LTM of flies fed high vs. low [Mg^2+^] diet that carried knockdown of *kibra*, *crb*, *wg*, or *Glut1* expression. Surprisingly, LTM defects of *crb* and *Glut1* knockdown flies were rescued by high [Mg^2+^] diet, whereas *kibra* and *wg* LTM defects were not (Figure 5C).

STM performance was not affected by knockdown of any of the target genes (Figure 5D) and diet-dependent memory enhancement was intact when these RNAi transgenes were not induced (Supplementary Figure 12C). To further support these findings, we again used FACS followed by RT-qPCR to examine αβ KC mRNA levels of Hippo pathway components and Sd target genes in flies fed low or high [Mg^2+^] diet. As predicted from the behavioral experiments, expression of *crb* and *Glut1*, but not *kibra* and *wg*, was elevated in flies fed high over low [Mg^2+^] diet (Supplementary Figure 13), consistent with *crb* and *Glut1* being key Hippo pathway target genes underlying the memory-enhancing effect of dietary Mg^2+^ supplement. However, individually overexpressing *crb* or *Glut1* in adult αβ KCs did not improve LTM (Figure 5E), showing that they are each necessary for high [Mg^2+^] diet to enhance LTM, but individually insufficient to enhance LTM. In contrast, LTM was significantly enhanced by adult αβ KC overexpression of *wg* (Figure 5E). None of these transgenes altered LTM when not induced (Supplementary Figure 12D). STM was also unaffected by overexpression of these Sd target genes (Supplementary Figure 14).

We lastly tested whether intact Hippo signaling was required for high [Mg^2+^] diet to rescue the LTM defect of flies expressing *crb* or *Glut1* RNAi in αβ KCs. Flies were constructed that co-expressed *crb* or *Glut1* RNAi with *hpo* RNAi in adult αβ KCs, and their LTM performance was compared between low and high [Mg^2+^] diets. In low [Mg^2+^] fed flies, LTM was impaired when *crb* or *Glut1* was knocked down, either with or without *hpo* knockdown (Figure 5F). However, high [Mg^2+^] diet no longer restored LTM in *crb* or *Glut1* knockdown flies that also lacked *hpo* (Figure 5F). Since *hpo* RNAi alone only impairs Mg^2+^-dependent LTM enhancement, and not baseline LTM (Figure 4, B and C), we conclude that dietary Mg^2+^ supplement enhances LTM by Hippo-dependent induction of Sd-regulated genes that include *crb* and *Glut1*.

## Discussion

In this study we demonstrate that upregulation of Hippo-dependent nuclear signaling underlies the memory-enhancing effects of dietary Mg^2+^ supplement. Mg^2+^ supplemented diet upregulates *Mer* and downregulates *hpo* in αβ KCs. More efficient Mer-directed membrane tethering of a smaller pool of available Hpo should disable Hpo’s ability to phosphorylate Wts, resulting in less Yki phosphorylation. Dephosphorylated Yki can then translocate to the nucleus, where it critically partners with Sd to enhance expression of key target genes. We discovered that LTM can be enhanced independently of diet by overexpressing *sd*, or releasing Sd from Tgi-mediated inhibition. Although Sd target genes *crb* and *Glut1*, in turn, support memory enhancement, their individual overproduction is insufficient, suggesting need for a larger Sd-directed transcriptional program, or additional independently regulated genes.

Recent work has shown that *crb* promotes axon pruning (*51*), which could indicate a role in long-term structural plasticity of memory relevant synaptic depression. It is also possible that *crb* induction provides negative-feedback on overactivation of Yki-dependent transcription. In epithelial cells, Crb recruits Ex to the apical membrane, thereby activating Hpo to inhibit Yki (*52*). However, we did not find a role for Ex in LTM, suggesting KC compartmentalization of Hpo signaling is independent of Ex. Our data instead demonstrate importance of Kibra, which can activate Hpo spatially distinct from and independent of Ex (*53*). Kibra provides synaptic scaffold for PKM-ζ that stabilizes synapses and promotes synaptic AMPA glutamate receptor trafficking in mammalian hippocampal synaptic plasticity and LTM (*54*, *55*). We note human polymorphism of Kibra (*56*), and patients with *CRB1* mutation-dependent retinopathy are associated with improved memory performance (*57*). Prior work has documented a role for KC-wide expression of *Glut1* in LTM formation and shown that neuronal glucose metabolism increases upon aversive LTM formation (*58*, *59*). However, neuronal glycolysis is dispensable for fly viability (*60*) and LTM was instead reported to require flux of KC glucose through the Pentose Phosphate pathway (PPP) (*58*), which produces ribose 5-phosphate, a critical precursor for nucleotide biosynthesis. Our analyses of Mg^2+^-dependent memory enhancement provide a unique perspective to the metabolic demands of LTM and support a role for the PPP and nucleotide synthesis. We also identified *CG2246*, which encodes fly PRPSAP (phosphoribosyl pyrophosphate synthetase associated protein), to be downregulated in αβ KCs by high [Mg^2+^] diet and for adult-restricted αβ KC *CG2246* knockdown to be memory enhancing, independently of diet. PRPSAP inhibits phosphoribosyl pyrophosphate synthetase (PRPS), which sits immediately downstream of the PPP, and converts ribose-5-phosphate into phosphoribosyl pyrophosphate (PRPP) (*61*). PRPSAP thereby controls the rate of ribose-5-phosphate use for nucleotide biosynthesis. We therefore propose that the memory-enhancing effect of dietary Mg^2+^ emerges from Hpo-dependent induction of Crb and Glut1, and accelerated conversion of imported glucose into nucleotide precursors for new gene expression to support structural synaptic remodeling (Supplementary Figure 15).

## Materials and Methods

### Experimental model and subject details

#### Drosophila strains

Flies were raised on standard cornmeal food under a 12 h/12 h light-dark cycle at 25 °C and 60% humidity, unless stated otherwise. Test and control flies for tubP-GAL80^ts^ experiments were raised at 18 °C. 5-day old mixed sex flies were used for the single-cell transcriptomics and RT-qPCR experiments. 1-7-day old mixed sex flies were used in the behavior experiments.

The wild-type strain was Canton-S. The GAL4 driver lines used were c739-GAL4 (*62*) and MB008B (*34*). The GAL80 line used was tubP-GAL80^ts^ (*43*). The UAS lines used were UAS-mCherry (*63*), UAS-*Mer*^RNAi^ (BDSC 34958), UAS-*jub*^RNAi^ (BDSC 41938), UAS-*E2f2*^RNAi^ (BDSC 27995), UAS-*SMSr*^RNAi^ (VDRC 39483), UAS-*Shc*^RNAi^ (BDSC 66961), UAS-*CG2926*^RNAi^ (BDSC 34941), UAS-*JIL-1*^RNAi^ (BDSC 57293), UAS-*c11.1*^RNAi^ (BDSC 61204), UAS-*SP1173*^RNAi^ (VDRC 109027), UAS-*bsk*^RNAi^ (BDSC 53310), UAS-*CG8485*^RNAi^ (BDSC 60083), UAS-*Crk*^RNAi^ (BDSC 55277), UAS-*CG15312*^RNAi^ (VDRC 101286), UAS-*babo*^RNAi^ (BDSC 25933), UAS-*CG10508*^RNAi^ (VDRC 102744), UAS-*CG6145*^RNAi^ (BDSC 57753), UAS-*krz*^RNAi^ (BDSC 33880), UAS-*CG2246*^RNAi^ (VDRC 110389), UAS-*RtGEF*^RNAi^ (VDRC 105093), UAS-*Prosbeta7*^RNAi^ (BDSC 34812), UAS-*hpo*^RNAi^ (BDSC 33614), UAS-*Spps*^RNAi^ (BDSC 62185), UAS-*kek2*^RNAi^ (BDSC 31874), UAS-*CG12219*^RNAi^ (BDSC 28000), UAS-*ksr*^RNAi^ (BDSC 32937), UAS-*Prp38*^RNAi^ (VDRC 110282), UAS-*ex*^RNAi^ (BDSC 28703), UAS-*sav*^RNAi^ (BDSC 28006), UAS-*wts*^RNAi^ (BDSC 27662), UAS-*mats*^RNAi^ (BDSC 34959), UAS-*Tgi*^RNAi^ (VDRC v31340), UAS-*yki*^RNAi^ (BDSC 34067), UAS-*sd*^RNAi^ (VDRC v101497), UAS-*Diap1*^RNAi^ (BDSC 33975), UAS-*CycE*^RNAi^ (BDSC 38902), UAS-*Myc*^RNAi^ (BDSC 43962), UAS-*kibra*^RNAi^ (BDSC 28683), UAS-*fj*^RNAi^ (BDSC 34323), UAS-*crb*^RNAi^ (BDSC 27697), UAS-*dally*^RNAi^ (BDSC 33952), UAS-*wg*^RNAi^ (BDSC 32994), UAS-*Glut1*^RNAi^ (VDRC 101365), UAS-*nerfin-1*^RNAi^ (BDSC 28324), UAS-*nerfin-1*^RNAi^ (VDRC 101631), UAS-Glut1 (*64*), UAS-Mer (BDSC 600495), UAS-*hpo* (BDSC 600509), UAS-*wts* (BDSC 600510), UAS-yki (BDSC 600526), UAS-*sd* (BDSC 9374), UAS-*kibra* (BDSC 600581), UAS-crb (BDSC 5544), UAS-*wg* (BDSC 5918), UAS-nerfin-1 (BDSC 67385). Control lines for TRiP RNAi lines are TRiP-attP2 (BDSC 36303) and TRiP-attP40 (BDSC 36304). Control lines for VDRC RNAi lines are v60100 and v60101.

## Method details

### [Mg^2+^] diet feeding

[Mg^2+^] feeding followed the protocol as described (*10*). 1-2 day old flies were raised in vials with Mg^2+^ supplemented food for 4 days before they were collected for experiments. Test groups were fed with high [Mg^2+^] (80mM) food while control groups were fed with low [Mg^2+^] (1mM) food. [Mg^2+^] food preparation was detailed in (*10*).

### Hpo/Sd inhibitor feeding

To investigate effects of small molecule inhibitors on memory, flies aged 1–2 days were cultured in vials containing food supplemented with specific compounds or DMSO as a control. Drug-containing food was prepared by adding the appropriate volume of a 10 mM stock solution to liquid food (e.g., 200 μL stock per 200 mL food for a 10 μM final concentration), followed by stirring at 600 rpm and incubation at 46 °C for one hour to ensure homogeneity. Food with additional concentrations was prepared similarly. Flies were fed the drug-supplemented food for 4 days before collection for downstream experiments.

For viability assays, 20 flies were maintained on 5 mL of food containing various drug concentrations for 7 days, after which the number of dead flies was recorded. The following inhibitors were used: XMU-MP-1 as Hpo inhibitor (*48*), and IK-930 as a Sd inhibitor (*46*).

### Brain dissociation and cell collection

Fly brain dissociation followed the protocol as described (*16*). For each sample, 10 female and 10 male brains were dissected in ice-cold DPBS (calcium and magnesium free, Gibco, 14190-086). Optic lobes were removed and the central brains were transferred immediately to 1 mL toxin-supplemented Schneider’s medium (tSM: Schneider’s medium (Gibco, 21720–001) containing 50 μM AP5 (Abcam, ab144482), 20 µM DNQX (6,7-dinitroquinoxaline-2,3-dione, ab14449), and 0.1 μM TTX (tetrodotoxin, ab120055)). Brains were then washed once with 1 mL tSM and incubated afterwards in 500 μL tSM containing 1 mg/ml papain (Sigma, P4762) and 1 mg/mL collagenase I (Sigma, C2674) for 20 min at room temperature. Next, brains were washed once more with tSM and subsequently triturated with flame-rounded 200 μL pipette tips. Dissociated brains were resuspended into 1 mL PBS + 0.01% BSA and filtered through a 10 µm CellTrix strainer (Sysmex, 04-0042-2314). Cell concentration was measured using a disposable Fuchs-Rosenthal hemocytometer (VWR, 631–1096) under a Leica DMIL LED Fluo microscope, which at the same time allowed a visualization of mCherry fluorescence in dissociated KCs. Cells were diluted to a concentration of up to 200 cells/μL in 400 μL PBS + 0.01% BSA solution. A typical preparation from 20 central brains yielded ∼500,000 single cells.

### 10x Genomics data processing

Single-cell libraries were made using the Chromium Single Cell 3ʹ v3 kit from 10x Genomics. Cells were loaded in accordance with 10x Genomics documentation. Samples were sequenced with 8 lanes of Illumina NovaSeq 6000 by Oxford Genomics Centre. Using CellRanger 4.1.0, barcodes were extracted and reads were aligned to a combination of the *Drosophila melanogaster* genome (R6.25) (*30*) and three reference sequences for mCherry and each split-GAL4 transgene of the flies that were used in this study (MB002B, MB077C, MB112C, MB434B). Flybase FB2018_06 gene names were used for the creation of the Digital Gene Expression (DGE) matrices.

### DGE matrix analysis with Seurat

Analysis of the DGE matrices with Seurat 4.0 (*65, 66*) followed the pipeline depicted in Figure 1B. In step 1 (basic quality control (QC)), cells that met one of the following criteria were removed:

<200 features

<800 UMIs

>40,000 UMIs

>15% mt-RNA

>15% ribosomal proteins

>60% ribosomal RNA

>10% TE

We obtained 51,232 cells from the raw data, and 9703 cells (18.94%) were filtered out in step 1, leaving 41,529 cells for the next step (Supplementary Figure 1G).

In step 2 (advanced QC), doublets were removed from each sample using DoubletFinder (*67*), following the tutorial on https://github.com/chris-mcginnis-ucsf/DoubletFinder. Overall, 3115 (7.50%) doublets were removed from the 41,529 cells, leaving 38,414 cells for the next step. Detailed cell numbers before and after each QC step are listed in Supplementary Figure 2C. In summary, we recovered 38,414 high quality singlets from the original 51,232 cells.

In step 3 (integration), condition-specific replicates (4 replicates for high [Mg^2+^] and 4 for low [Mg^2+^]) were merged separately using Seurat function “merge”. Next, the two merged groups were processed through Seurat function “SCTransform”, which packages normalization, scaling and variable feature identification (*12*). Also, during normalization, confounding sources of variation such as mitochondrial percentage, rRNA percentage, and sex-specific expression were regressed out. Finally, the two groups were integrated through Seurat function “IntegrateData” using canonical correlation analysis (CCA) with 2,000 highly variable features selected before (*13*).

In step 4 (clustering), dimension reduction of the integrated dataset (from 2,000 to 30) was carried out using principle component analysis (PCA). Further, uniform manifold approximation and projection (UMAP) was performed to reduce the dimension to 2D in order to visualize the cells (*15*). Next, clusters were identified through a modularity optimization algorithm based on a shared nearest neighbor measurement, using a resolution of 1.0 (*14*).

In step 5 (cell typing), clusters were annotated by their expression of known marker genes: *crb* for αβ Kenyon cells (KCs), *ab* for γ KCs, *CG8641* for α′β′ KCs, *ple* for dopaminergic neurons, *alrm* for astrocyte, *Poxn* for ellipsoid body and *wtrw* for ensheathing glia (Figure 1C).

In step 6 (differential expression (DE) analysis), three different methods were used to compare the expression of genes for each typed cluster between high and low [Mg^2+^] conditions.

First, we took advantage of the toolset “Muscat”, which adapted DE algorithms edgeR (*68*) and limma (*69*) for the multi-sample, multi-condition single-cell transcriptomics data (*70*). Running process followed the tutorial (https://github.com/HelenaLC/muscat/blob/master/vignettes/analysis.Rmd), with a threshold of 0.2 set for log2FC, 0.05 set for adjusted p-val, and “edgeR” set for method.

Second, a battery of alternative tests was performed through the Seurat function “FindMarkers()” following the Seurat DE tutorial (https://satijalab.org/seurat/articles/de_vignette.html), and their respective results were integrated afterwards. We named this method “Cocktail” for convenience. Those tests include “wilcox” (Wilcoxon rank sum test), “bimod” (likelihood-ratio test, (*71*)), “poisson” (Likelihood ratio test assuming an underlying poisson distribution), “negbinom” (Likelihood ratio test assuming an underlying negative binomial distribution), “LR” (Uses a logistic regression framework to determine differentially expressed genes, constructs a logistic regression model predicting), “MAST” (GLM-framework that treats cellular detection rate as a covariate, (*72*)), “DEseq2” (DE based on a model using the negative binomial distribution, (*73*)). nUMI and sex-specific gene expression were added as latent variables for poisson, negbinom, LR and MAST tests. The results from those tests were assembled, and the genes with a |log2FC| >0.2 and with either an FDR adjusted p-val<0.01 from at least one test, or p-val<0.05 reported from at least 3 different tests were collected as DEGs.

In a third method, which we named as “DirectE”, we used the Seurat function “AverageExpression” to calculate a pseudo-bulk expression level based on the raw RNA counts for each gene in any specific cluster. We then compared directly the expression levels of each gene from the 4 high [Mg^2+^] samples with those from the 4 low [Mg^2+^] samples using Student’s t-test, for each typed cluster. A criterion of FDR adjusted p-val<0.05 and |log2FC|>0.2 was set for significantly different expression levels.

Finally, differentially expressed genes between the high [Mg^2+^] and low [Mg^2+^] conditions from the above 3 methods were collected together, and their overlap profiles were generated as Venn diagrams.

### Fluorescence-activated cell sorting (FACS) and quantitative real-time PCR (qPCR)

Flies expressing mCherry in αβ KCs, driven by MB008B GAL4, were fed with high or low [Mg^2+^] supplemented food for 4 days before dissection. Six independent samples were prepared for each different treatment or genotype. For each sample, central brains were dissected and dissociated as above. Filtered brain cells were then sorted with a FACS Aria III (BD Biosciences) and mCherry+ cells were collected. Cell lysis, reverse transcription and first-strand cDNA synthesis were performed using the SuperScript IV CellsDirect cDNA Synthesis Kit (Invitrogen, 11750150) following the manufacturer’s instructions. Obtained cDNA was pre-amplified with the KAPA HiFi HotStart ReadyMix (Kapa Biosystems, KK2611). Each 50 μL reaction contained ∼1 μg of cDNA template, 25 μL of 2X KAPA HiFi HotStart ReadyMix, and 20 μL of primer mix (1 μM). Pre-amplification PCR protocol was as follows: 95 °C, 3m; 18X [98 °C, 20 s; 60 °C, 15 s; 72 °C, 15 s]; 72 °C, 15 s. Pre-amplified cDNA was purified with the QIAquick PCR Purification Kit (Qiagen, 28106). Quantitative PCR was performed in triplicate for each cDNA sample on a LightCycler 480 Instrument (Roche) using SYBR Green I Master Mix (Roche, 04707516001). Each 10 μL reaction contained 1 μL of pre-amplified cDNA, 0.4 μM of each primer, and 5 μL of SYBR Green I Master Mix. Cycles were as follows: 95 °C, 10m; 45X [95 °C, 10 s; 60 °C, 20 s; Fluorescence acquisition; 72 °C, 10 s]. Melting curves were analyzed after amplification, and amplicons were visualized by agarose gel electrophoresis to confirm primer specificity. Relative transcript levels were calculated by the 2-ΔΔCt method (*74*), using SdhA as reference gene (*16, 75, 76*). Primers are detailed in table S1.

### uex::HA imaging quantification in αβ Kenyon cells

*uex*::HA flies (*10*) were crossed to c739-GAL4; UAS-mCherry to label αβ KCs with mCherry. Flies were fed with either 1mM (low [Mg^2+^]) or 80mM (high [Mg^2+^]) for 4 days before dissection.

Immunostaining was performed as described previously (*77*). Briefly, brains were dissected in ice-cold PBS and fixed in 4% paraformaldehyde in PBS for 20 min at room temperature. Samples were washed twice briefly and three times for 20 min in 0.5% PBT (PBS containing 0.5% Triton X-100), blocked for 30 min in 5% normal goat serum in PBT, and incubated with primary and secondary antibodies for 48 h each at 4 °C with gentle rotation (50 rpm). The primary antibody was rat anti-HA (1:250; Merck, 11867423001), and the secondary antibody was Alexa Fluor 488-conjugated goat anti-rat (1:250; Invitrogen, A11006). Brains were washed twice briefly and three times for 20 min in PBT before and after secondary antibody incubation. DAPI (1:1000; Thermo Scientific, 62248) was included during the second 20-min wash after secondary antibody incubation. Brains were mounted in Vectashield (Vector Laboratories, H1000).

Images were acquired on a Zeiss LSM 900 confocal microscope equipped with an Airyscan detector using a Plan-Apochromat 63×/1.4 NA oil objective. Three-channel z-stacks spanning the αβ KC cell-body layer were collected at 2186 × 2186 pixel resolution with 1-µm z-steps. Channels comprised c739-GAL4-driven mCherry fluorescence marking αβ KC cell bodies, anti-HA immunofluorescence reporting *uex*::HA, and DAPI labeling nuclei. Images were processed in Fiji (*78*).

*uex*::HA enrichment in αβ KCs was quantified using a custom Python pipeline adapted from (*79, 80*). Native CZI files were read using aicsimageio and aicspylibczi, and each optical section was analyzed independently. Cell bodies were segmented from the mCherry channel and nuclei from the DAPI channel using the Cellpose-SAM generalist model (*81*). Putative αβ KCs were identified by calculating the mean mCherry intensity of each segmented cell and classifying cells using an Otsu threshold (scikit-image; (*82*)) applied to the per-cell mCherry intensity distribution. Cells above the threshold were classified as mCherry-positive αβ KCs, whereas all remaining cells constituted the baseline population.

For each segmented cell, *uex*::HA abundance was measured as the mean anti-HA fluorescence intensity across the cell body. The αβ KC signal was defined as the mean anti-HA intensity of all mCherry-positive cells. Baseline fluorescence was measured from mCherry-negative cells using an equivalent whole-cell compartment generated by expanding each DAPI-labeled nucleus with a perinuclear ring whose width equaled the difference between the median cell-body radius and the median nuclear radius for that optical section (implemented using expand_labels in scikit-image), while excluding mCherry-positive pixels. This procedure matched the signal and baseline measurement compartments in size.

Per-brain normalized *uex*::HA values were compared between flies fed with low [Mg^2+^] food (1 mM; n = 10 brains) and high [Mg^2+^] food (80 mM; n = 12 brains) using a two-sided Mann– Whitney U test implemented in SciPy (*83*). Statistical significance was defined as *P* < 0.05.

### Behavioral experiments

Behavioral experiments were carried out using the T-maze apparatus (*84*), and the protocols are described in detail in (*10*). Flies used were 1-7 day old of mixed sex. Odors were 4-methylcyclohexanol (MCH) and 3-octanol (OCT), diluted ∼1:10^3^ in mineral oil. All experiments were conducted at 23 °C and 55%-65% relative humidity. For experiments using tubP-GAL80^ts^, flies were collected from 18 °C incubator after eclosion and put into 18 °C (for uninduced group) or 31 °C (for induced group) incubator for 3-4 days before they are aliquoted for behavior experiments; 31 °C is 1 °C above the restrictive temperature, to ensure complete inactivation of GAL80^ts^.

Appetitive memory assay was performed as described (*20*, *23*). Groups of ∼100 flies were starved for 21-23 h before training. During training, flies were loaded into a tube coated inside with paper dried from water soaking, and exposed to CS-odor for 2 min. This was followed by 30 s of clean air. Flies were then transferred to another tube coated with paper dried from sugar soaking, and were exposed to the CS+ odor for 2 min. After training, flies were either transported to the choice point for testing of immediate memory (STM), or they were fed for 40-60 min on standard food and transferred back to the starvation vials for 23 h until testing for LTM. Performance Index was calculated as the number of flies in the CS+ arm minus the number in the CS-arm, divided by the total number of flies. MCH and OCT, were alternately used as CS+ or CS-and a single sample, or n, represents the average performance score from two reciprocally trained groups. [Mg^2+^] food feeding was performed as above before starvation. Sensory acuity tests were performed as described (*10, 85–87*). To test olfactory acuity, untrained flies were given 2 min to choose between a diluted odor as used in training and air bubbled through mineral oil in the T-maze. An Avoidance Index was calculated as the number of flies in the air arm minus the number in the odor arm, divided by the total number of flies. To assess sugar acuity, starved flies were given 2 min to choose between an arm of the T-maze containing a dried sugar paper and the other containing a dried ‘water’ filter paper. Both papers were prepared as in the appetitive memory assays. A Preference Index was calculated as the number of flies in the sugar arm minus that in the other arm, divided by the total number of flies. Light was turned on and air flow was allowed to run through the testing tubes during sugar preference assay.

### Quantification and statistical analyses

Behavior data were analyzed using Excel and Prism 6. Unpaired two-tailed t-tests were used for comparing two groups, and one-way ANOVA followed by a Tukey’s post-hoc test was used for comparing multiple groups. Definition of statistical significance is set at *P* < 0.05.

## Data, code, and materials availability

Behavior data from T-maze assays are deposited in the Dryad Digital Repository (doi:10.5061/dryad.k0p2ngfqg). The single-cell RNA-sequencing data generated in this study have been deposited in the NCBI Sequence Read Archive (SRA) under BioProject accession PRJNA1514305. Analysis code is available at https://github.com/rdcwuyy/memory-enhancement-mg/. All other data are available in the main text or the supplementary materials.

## Acknowledgments

We thank V. Tsioligka and R. Hedley at the Flow Cytometry Facility of the Sir William Dunn School of Pathology, Oxford for their help with FACS. We thank D. Agarwal for support with Seurat analyses and A. Cook and I. Kapoor for comments on the manuscript.

We thank L. Partridge and her lab for flies. This work was supported by a Wellcome Principal Research Fellowship (200846), a Wellcome Discovery Award (225192), a European Research Council Advanced Grant (789274), and Wellcome Collaborative Awards (203261 and 209235) to SW.

## Author contributions

Conceptualization: SW, YW; Methodology: SW, YW, CT; Investigation: SW, YW; Formal analysis: SW, YW, CT; Visualization: YW; Funding acquisition: SW; Supervision: SW; Writing – original draft: SW, YW; Writing – review & editing: SW, YW, CT.

