## Supplementary Information for "Dietary magnesium supplement enhances memory through Hippo signaling"

### Supplementary Figures

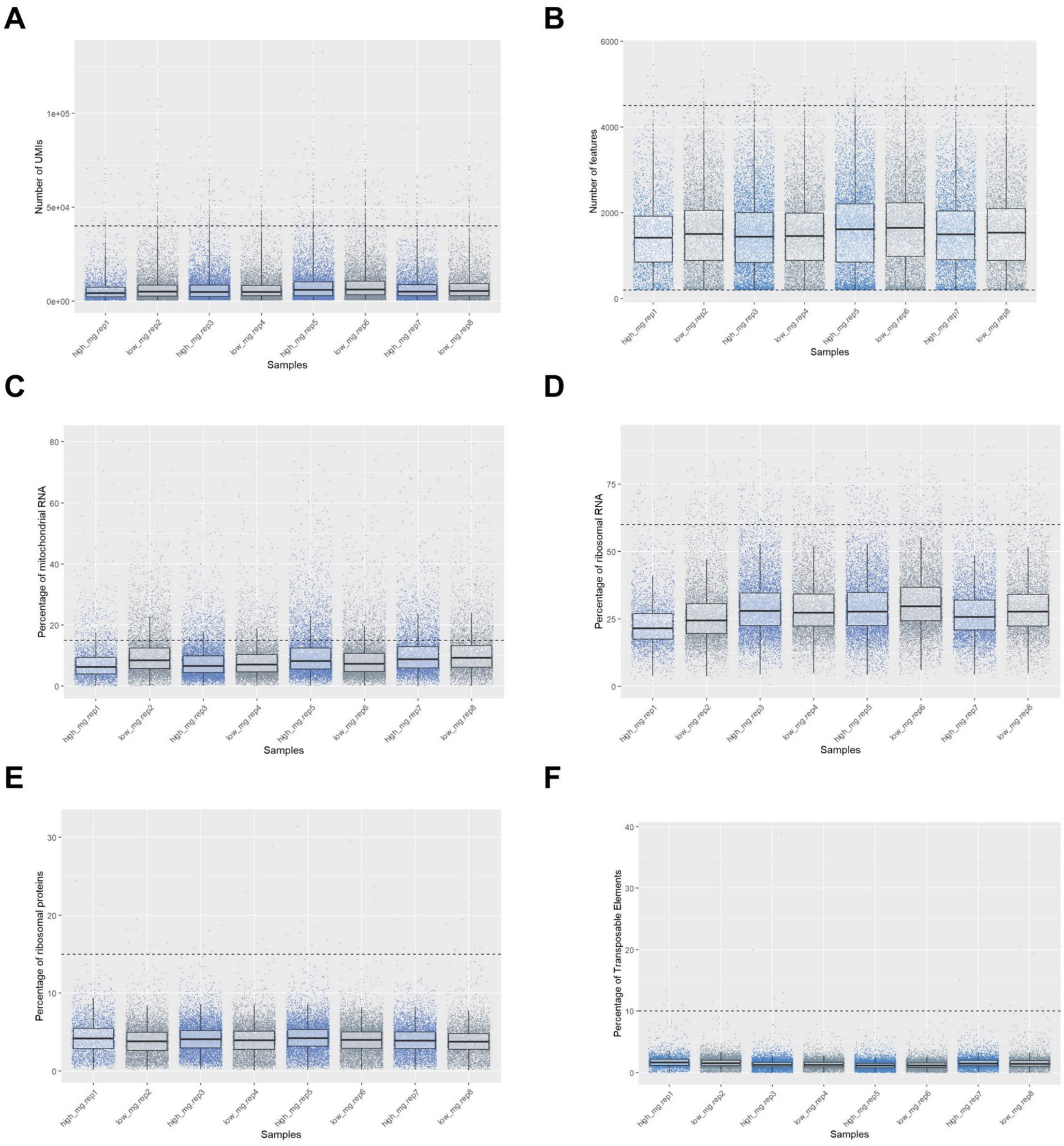


**G**

| Sample | Treatment | Raw cell number | Number (percent) of cells removed in basic QC | Number of Cells after basic QC |
|---|---|---|---|---|
| 1 | high $Mg^{2+}$ | 3437 | 517 (15.04%) | 2920 |
| 2 | low $Mg^{2+}$ | 6332 | 1398 (22.08%) | 4934 |
| 3 | high $Mg^{2+}$ | 8426 | 1185 (14.06%) | 7241 |
| 4 | low $Mg^{2+}$ | 6307 | 987 (15.65%) | 5320 |
| 5 | high $Mg^{2+}$ | 8053 | 1735 (21.54%) | 6318 |
| 6 | low $Mg^{2+}$ | 7651 | 1322 (17.28%) | 6329 |
| 7 | high $Mg^{2+}$ | 6151 | 1368 (22.24%) | 4783 |
| 8 | low $Mg^{2+}$ | 4875 | 1191 (24.43%) | 3684 |
| **Total** | | **51232** | **9703 (18.94%)** | **41529** |

**Supplementary Figure 1. Basic quality control.** (**A** to **F**) Low-quality cells were filtered out according to 7 criteria; each dot represents one cell and the dashed lines represent the thresholds (see materials and methods). (**G**) Cell numbers before and after basic quality control.

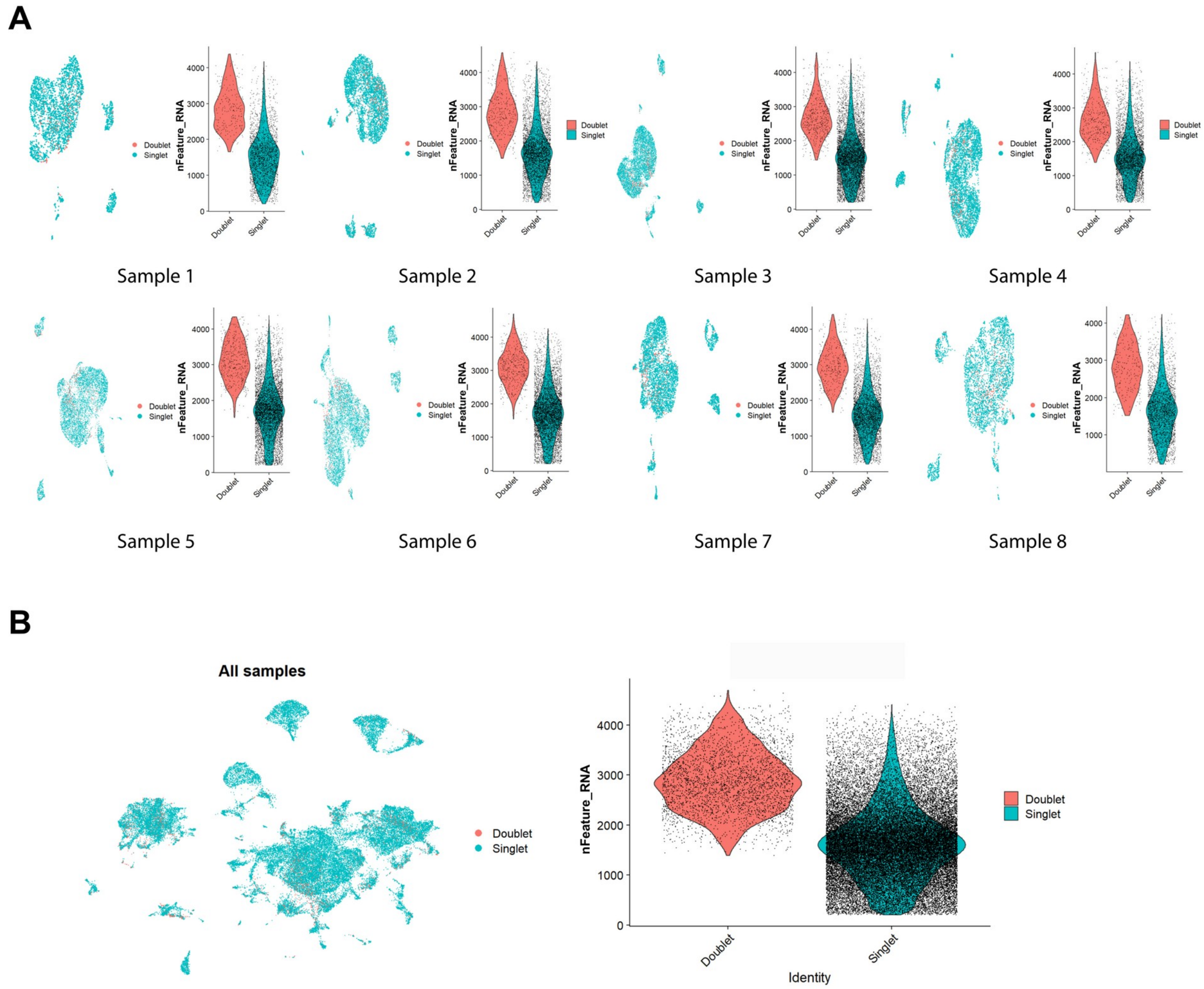


C

| Sample | Treatment | Raw cell number | Number of Cells after basic QC | Number (percent) of cells removed as doublets | Number of cells after doublet removal |
|---|---|---|---|---|---|
| 1 | high $Mg^{2+}$ | 3437 | 2920 | 219 (7.5%) | 2701 |
| 2 | low $Mg^{2+}$ | 6332 | 4934 | 370 (7.5%) | 4564 |
| 3 | high $Mg^{2+}$ | 8426 | 7241 | 543 (7.5%) | 6698 |
| 4 | low $Mg^{2+}$ | 6307 | 5320 | 399 (7.5%) | 4921 |
| 5 | high $Mg^{2+}$ | 8053 | 6318 | 474 (7.5%) | 5844 |
| 6 | low $Mg^{2+}$ | 7651 | 6329 | 475 (7.51%) | 5854 |
| 7 | high $Mg^{2+}$ | 6151 | 4783 | 359 (7.51%) | 4424 |
| 8 | low $Mg^{2+}$ | 4875 | 3684 | 276 (7.49%) | 3408 |
| **Total** | | **51232** | **41529** | **3115 (7.50%)** | **38414** |

**Supplementary Figure 2. Advanced quality control: removal of doublets.** (**A**) Sample-wise distribution of doublets. For each sample, a UMAP visualization of the distribution of doublets and singlets (left) and a violin plot showing the number of detected genes (nFeature_RNA) for doublets and singlets (right). (**B**) Distribution and gene counts for doublets across all samples. (**C**) Cell numbers before and after doublet removal.

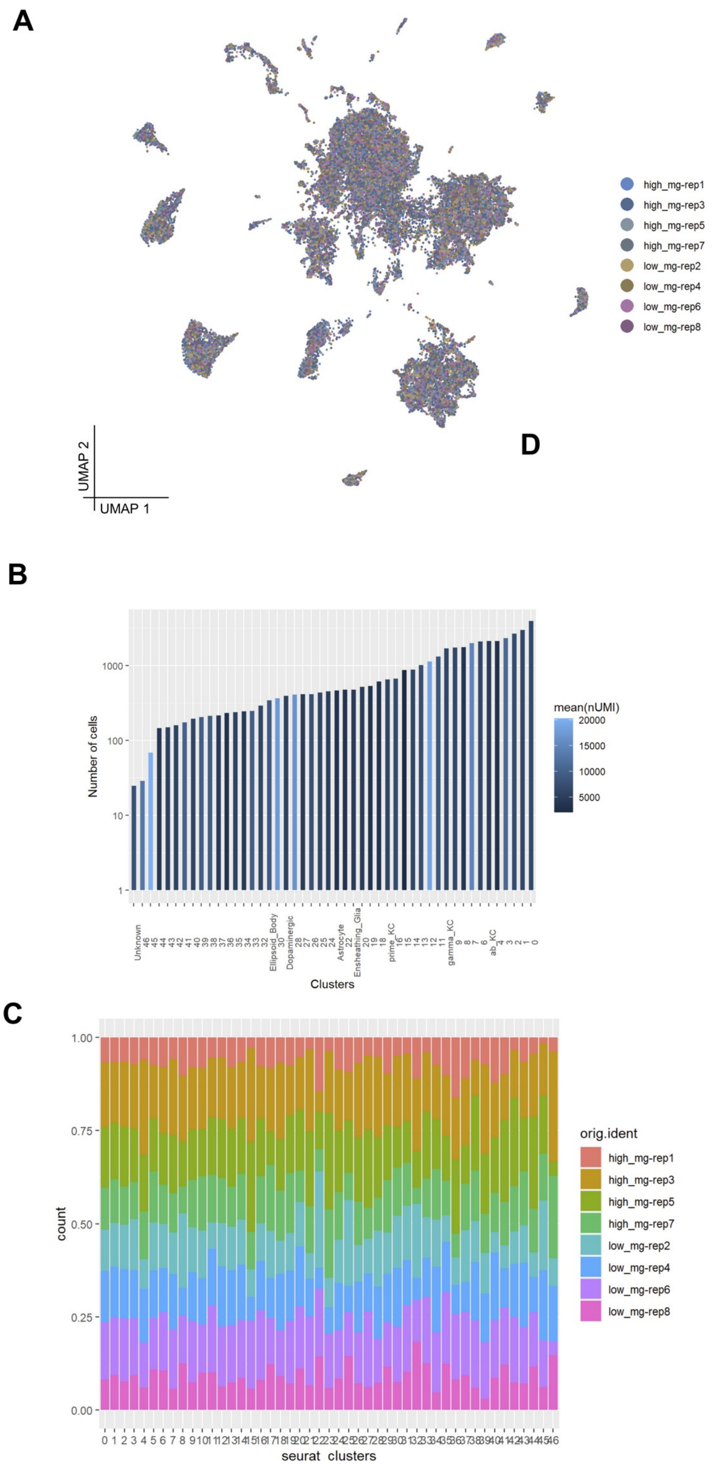


**Supplementary Figure 3. Composition of the cell clusters.** (**A**) Contribution of cells from the 8 samples to each cluster in the UMAP plot. (**B**) Number of cells in each identified cell cluster. (**C**) Percentage of counts contributed by the 8 samples to each cluster.

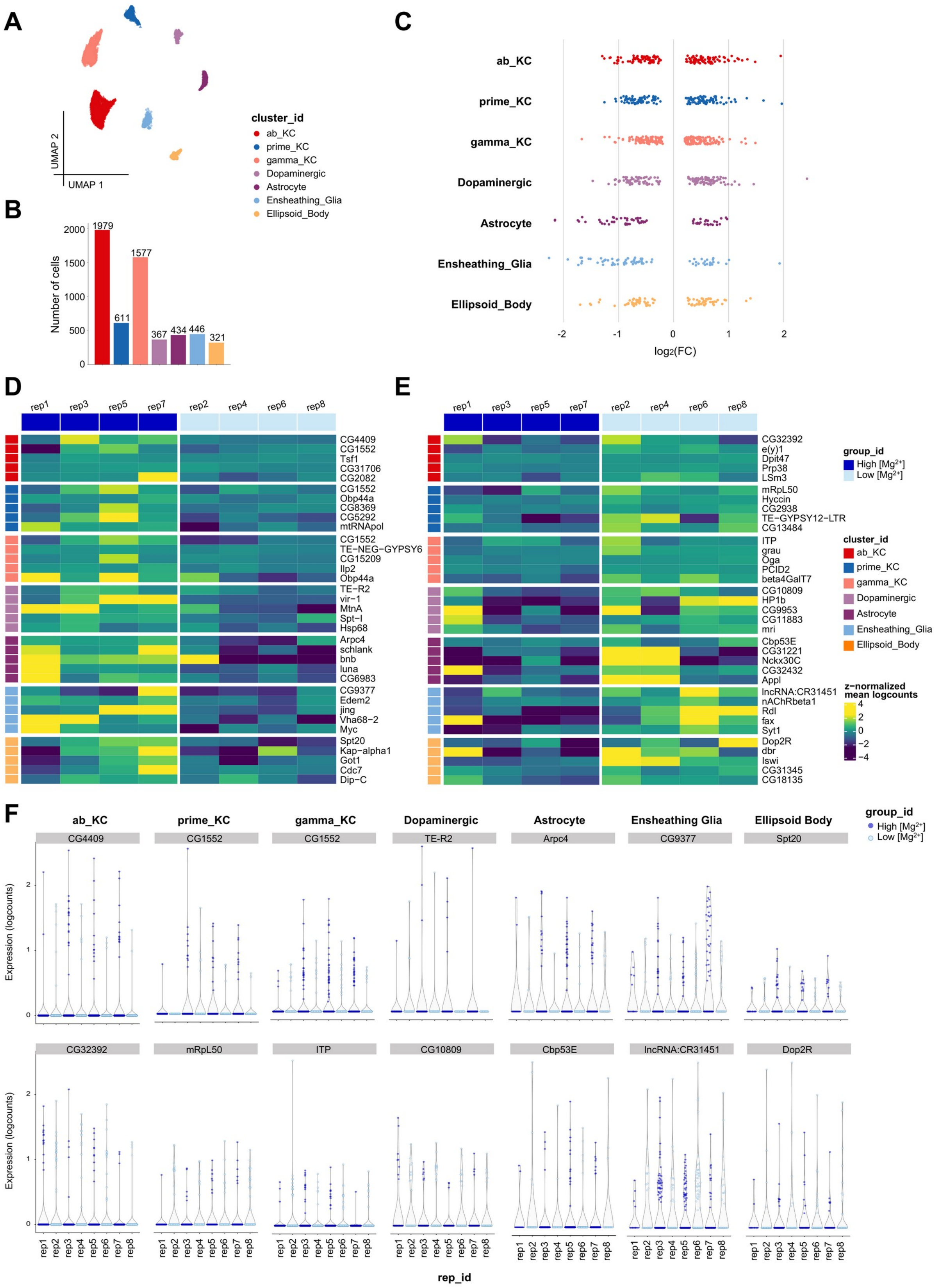
A
B
C
D
E
F
UMAP 2
UMAP 1
cluster_id
ab_KC
prime_KC
gamma_KC
Dopaminergic
Astrocyte
Ensheathing_Glia
Ellipsoid_Body
Number of cells
1979
611
1577
367
434
446
321
log2(FC)
group_id
High [Mg2+]
Low [Mg2+]
rep1
rep3
rep5
rep7
rep2
rep4
rep6
rep8
CG4409
CG1552
Tsf1
CG31706
CG2082
Obp44a
CG8369
CG5292
mtRNApol
TE-NEG-GYPSY6
CG15209
Ilp2
TE-R2
vir-1
MtnA
Spt-I
Hsp68
Arpc4
schlank
bnb
luna
CG6983
CG9377
Edem2
jing
Vha68-2
Myc
Spt20
Kap-alpha1
Got1
Cdc7
Dip-C
CG32392
e(y)1
Dpit47
Prp38
LSm3
mRpL50
Hyccin
CG2938
TE-GYPSY12-LTR
CG13484
ITP
grau
Oga
PCID2
beta4GalT7
CG10809
HP1b
CG9953
CG11883
mri
Cbp53E
CG31221
Nckx30C
CG32432
Appl
lncRNA:CR31451
nAChRbeta1
Rdl
fax
Syt1
Dop2R
dbr
Iswi
CG31345
CG18135
z-normalized mean logcounts
Ensheathing Glia
Ellipsoid Body
Expression (logcounts)
rep_id

20

**Supplementary Figure 4. Differentially expressed genes in the seven major cell types.** (A) UMAP visualization of the seven major cell types. (B) Number of cells in each of the seven major cell types. (C) Dot plot showing the distribution of $\log_2$ fold changes of the DEGs detected by Muscat in each of the seven major cell types ($|\log_2 FC| > 0.2$); each dot represents one DEG. (D and E) Heatmaps showing the top 5 up-regulated (D) and top 5 down-regulated (E) DEGs of each of the seven major cell types. In each heatmap the four high [$Mg^{2+}$] samples are displayed on the left and the four low [$Mg^{2+}$] samples on the right. The color of each cell codes the z-normalized mean log(counts) of the corresponding gene. (F) Violin plots showing the expression level, in individual cells of each sample, of the most up-regulated (upper row) and most down-regulated (lower row) DEG of each major cell type.

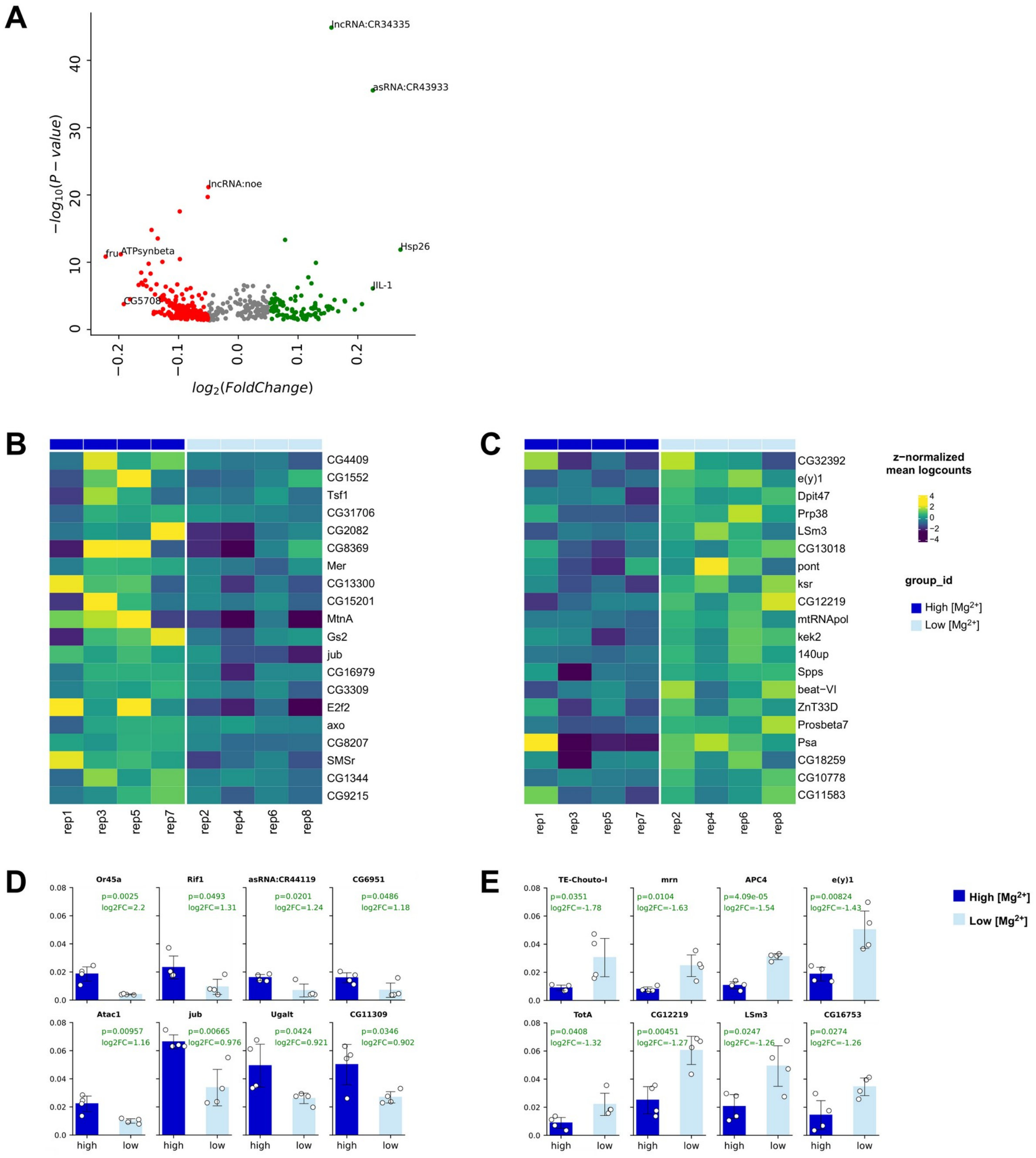


**Supplementary Figure 5. Outputs of the three individual differential expression methods for αβ Kenyon cells.** (**A**) Volcano plot showing the DEGs identified by the Cocktail DE method. (**B** and **C**) Heatmaps showing the top 10 up-regulated (B) and top 10 down-regulated (C) DEGs identified by the Muscat DE method. The color of each cell codes the z-normalized mean log(counts) of the corresponding gene. (**D** and **E**) Bar plots showing the top 8 up-regulated (D) and top 8 down-regulated (E) DEGs identified by the DirectE DE method. The $log_2$FC and *P* value are labeled on each bar plot.

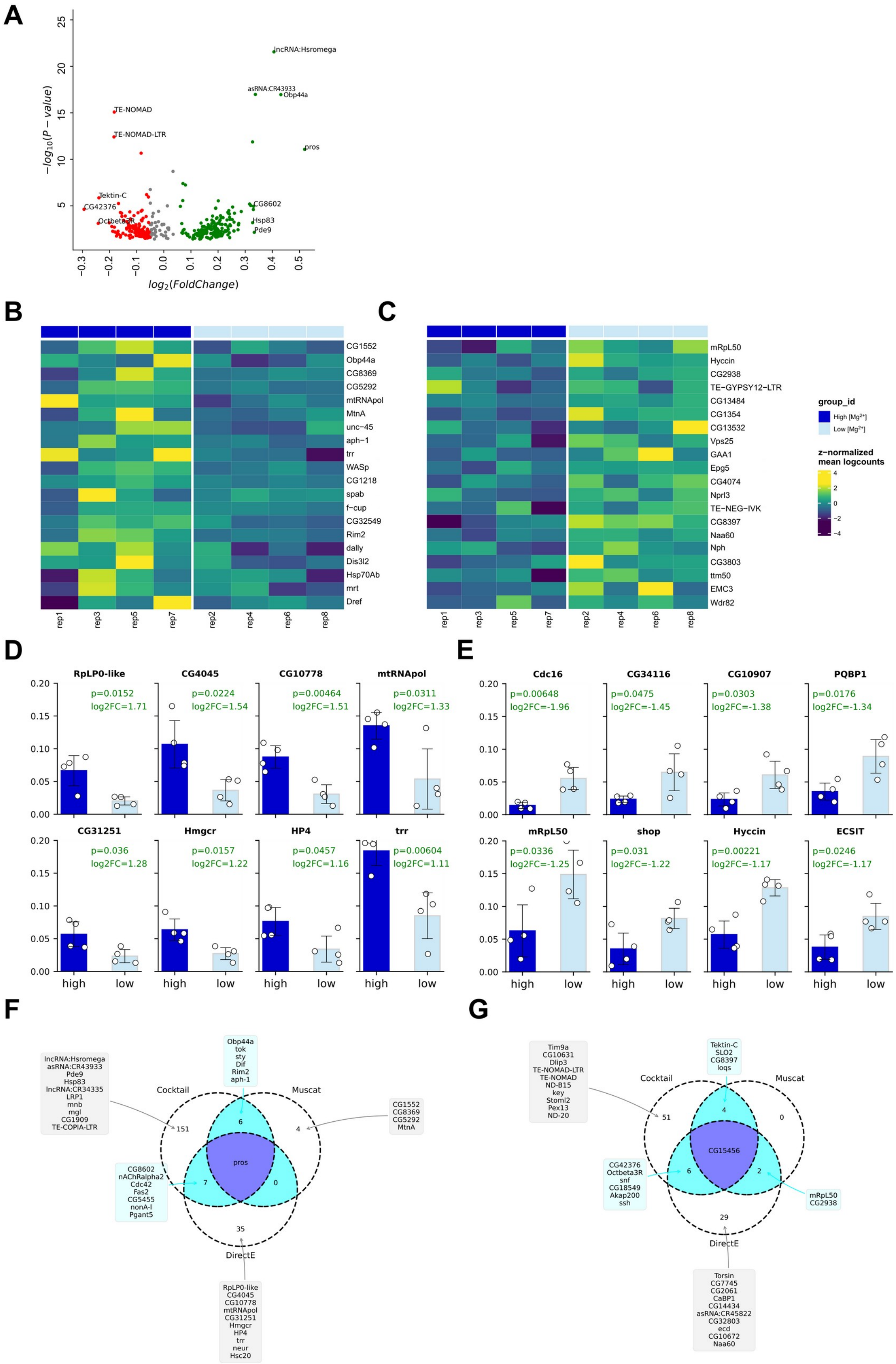
A
lncRNA:Hsromega
asRNA:CR43933
Obp44a
TE-NOMAD
TE-NOMAD-LTR
pros
Tektin-C
CG42376
Octbeta3R
CG8602
Hsp83
Pde9
−log10(P − value)
log2(FoldChange)
B
CG1552
Obp44a
CG8369
CG5292
mtRNApol
MtnA
unc−45
aph−1
trr
WASp
CG1218
spab
f−cup
CG32549
Rim2
dally
Dis3l2
Hsp70Ab
mrt
Dref
rep1
rep3
rep5
rep7
rep2
rep4
rep6
rep8
C
mRpL50
Hyccin
CG2938
TE−GYPSY12−LTR
CG13484
CG1354
CG13532
Vps25
GAA1
Epg5
CG4074
Nprl3
TE−NEG−IVK
CG8397
Naa60
Nph
CG3803
ttm50
EMC3
Wdr82
group_id
High [Mg2+]
Low [Mg2+]
z−normalized
mean logcounts
D
RpLP0-like
p=0.0152
log2FC=1.71
CG4045
p=0.0224
log2FC=1.54
CG10778
p=0.00464
log2FC=1.51
mtRNApol
p=0.0311
log2FC=1.33
CG31251
p=0.036
log2FC=1.28
Hmgcr
p=0.0157
log2FC=1.22
HP4
p=0.0457
log2FC=1.16
trr
p=0.00604
log2FC=1.11
high
low
E
Cdc16
p=0.00648
log2FC=-1.96
CG34116
p=0.0475
log2FC=-1.45
CG10907
p=0.0303
log2FC=-1.38
PQBP1
p=0.0176
log2FC=-1.34
mRpL50
p=0.0336
log2FC=-1.25
shop
p=0.031
log2FC=-1.22
Hyccin
p=0.00221
log2FC=-1.17
ECSIT
p=0.0246
log2FC=-1.17
F
Cocktail
Muscat
DirectE
lncRNA:Hsromega asRNA:CR43933 Pde9 Hsp83 lncRNA:CR34335 LRP1 mnb mgl CG1909 TE-COPIA-LTR
Obp44a tok sty Dif Rim2 aph-1
CG1552 CG8369 CG5292 MtnA
CG8602 nAChRalpha2 Cdc42 Fas2 CG5455 nonA-l Pgant5
RpLP0-like CG4045 CG10778 mtRNApol CG31251 Hmgcr HP4 trr neur Hsc20
151
6
4
pros
7
0
35
G
Tim9a CG10631 Dlip3 TE-NOMAD-LTR TE-NOMAD ND-B15 key Stoml2 Pex13 ND-20
Tektin-C SLO2 CG8397 loqs
Cocktail
Muscat
51
4
0
CG15456
CG42376 Octbeta3R snf CG18549 Akap200 ssh
6
2
mRpL50 CG2938
29
DirectE
Torsin CG7745 CG2061 CaBP1 CG14434 asRNA:CR45822 CG32803 ecd CG10672 Naa60

**Supplementary Figure 6. DEGs of α′β′ Kenyon cells detected by the combinatory DE analysis.** (**A**) Volcano plot showing the DEGs identified by the Cocktail DE method. (**B** and **C**) Heatmaps showing the top 10 up-regulated (B) and top 10 down-regulated (C) DEGs identified by the Muscat DE method. The color of each cell codes the z-normalized mean log(counts) of the corresponding gene. (**D** and **E**) Bar plots showing the top 8 up-regulated (D) and top 8 down-regulated (E) DEGs identified by the DirectE DE method. The $log_2FC$ and *P* value are labeled on each bar plot. (**F** and **G**) Venn diagrams showing the integrated DE results from the three methods: (F) up-regulated DEGs, (G) down-regulated DEGs.

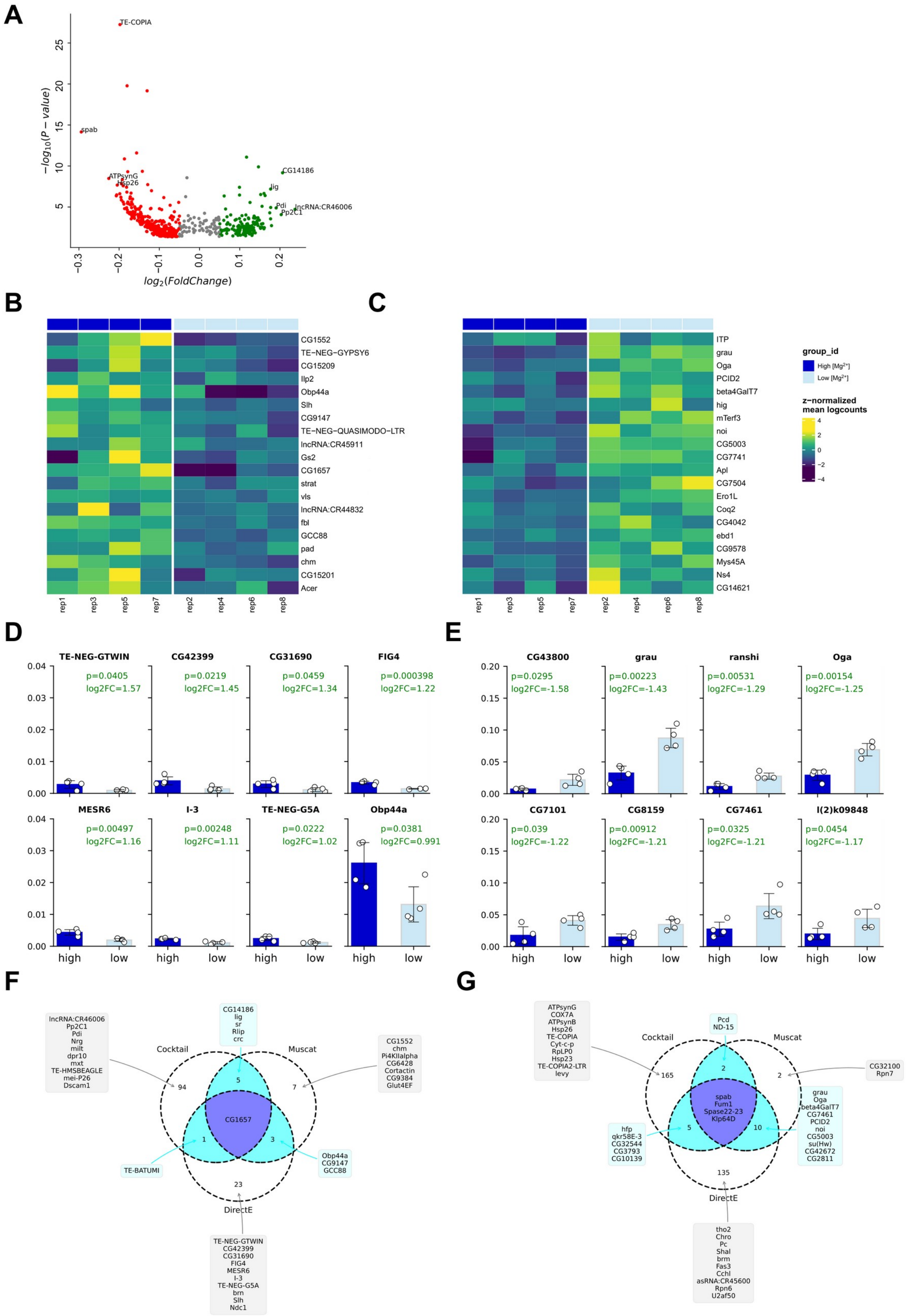
A
TE-COPIA
spab
ATPsynG
Hsp26
CG14186
lig
Pdi
lncRNA:CR46006
Pp2C1
$-log_{10}(P-value)$
$log_2(FoldChange)$
B
CG1552
TE−NEG−GYPSY6
CG15209
Ilp2
Obp44a
Slh
CG9147
TE−NEG−QUASIMODO−LTR
lncRNA:CR45911
Gs2
CG1657
strat
vls
lncRNA:CR44832
fbl
GCC88
pad
chm
CG15201
Acer
rep1 rep3 rep5 rep7 rep2 rep4 rep6 rep8
C
ITP
grau
Oga
PCID2
beta4GalT7
hig
mTerf3
noi
CG5003
CG7741
Apl
CG7504
Ero1L
Coq2
CG4042
ebd1
CG9578
Mys45A
Ns4
CG14621
rep1 rep3 rep5 rep7 rep2 rep4 rep6 rep8
group_id
High [Mg2+]
Low [Mg2+]
z−normalized mean logcounts
D
TE-NEG-GTWIN p=0.0405 log2FC=1.57
CG42399 p=0.0219 log2FC=1.45
CG31690 p=0.0459 log2FC=1.34
FIG4 p=0.000398 log2FC=1.22
MESR6 p=0.00497 log2FC=1.16
I-3 p=0.00248 log2FC=1.11
TE-NEG-G5A p=0.0222 log2FC=1.02
Obp44a p=0.0381 log2FC=0.991
high low
E
CG43800 p=0.0295 log2FC=-1.58
grau p=0.00223 log2FC=-1.43
ranshi p=0.00531 log2FC=-1.29
Oga p=0.00154 log2FC=-1.25
CG7101 p=0.039 log2FC=-1.22
CG8159 p=0.00912 log2FC=-1.21
CG7461 p=0.0325 log2FC=-1.21
l(2)k09848 p=0.0454 log2FC=-1.17
high low
F
Cocktail
Muscat
DirectE
CG14186 lig sr Rlip crc
lncRNA:CR46006 Pp2C1 Pdi Nrg milt dpr10 mxt TE-HMSBEAGLE mei-P26 Dscam1
CG1552 chm Pi4KIIalpha CG6428 Cortactin CG9384 Glut4EF
CG1657
TE-BATUMI
Obp44a CG9147 GCC88
TE-NEG-GTWIN CG42399 CG31690 FIG4 MESR6 I-3 TE-NEG-G5A brn Slh Ndc1
G
ATPsynG COX7A ATPsynB Hsp26 TE-COPIA Cyt-c-p RpLP0 Hsp23 TE-COPIA2-LTR levy
Pcd ND-15
CG32100 Rpn7
spab Fum1 Spase22-23 Klp64D
hfp qkr58E-3 CG32544 CG3793 CG10139
grau Oga beta4GalT7 CG7461 PCID2 noi CG5003 su(Hw) CG42672 CG2811
tho2 Chro Pc Shal brm Fas3 CCHa1 asRNA:CR45600 Rpn6 U2af50

48

**Supplementary Figure 7. DEGs of γ Kenyon cells detected by the combinatory DE analysis.** (**A**) Volcano plot showing the DEGs identified by the Cocktail DE method. (**B** and **C**) Heatmaps showing the top 10 up-regulated (B) and top 10 down-regulated (C) DEGs identified by the Muscat DE method. The color of each cell codes the z-normalized mean log(counts) of the corresponding gene. (**D** and **E**) Bar plots showing the top 8 up-regulated (D) and top 8 down-regulated (E) DEGs identified by the DirectE DE method. The $log_2FC$ and *P* value are labeled on each bar plot. (**F** and **G**) Venn diagrams showing the integrated DE results from the three methods: (F) up-regulated DEGs, (G) down-regulated DEGs.

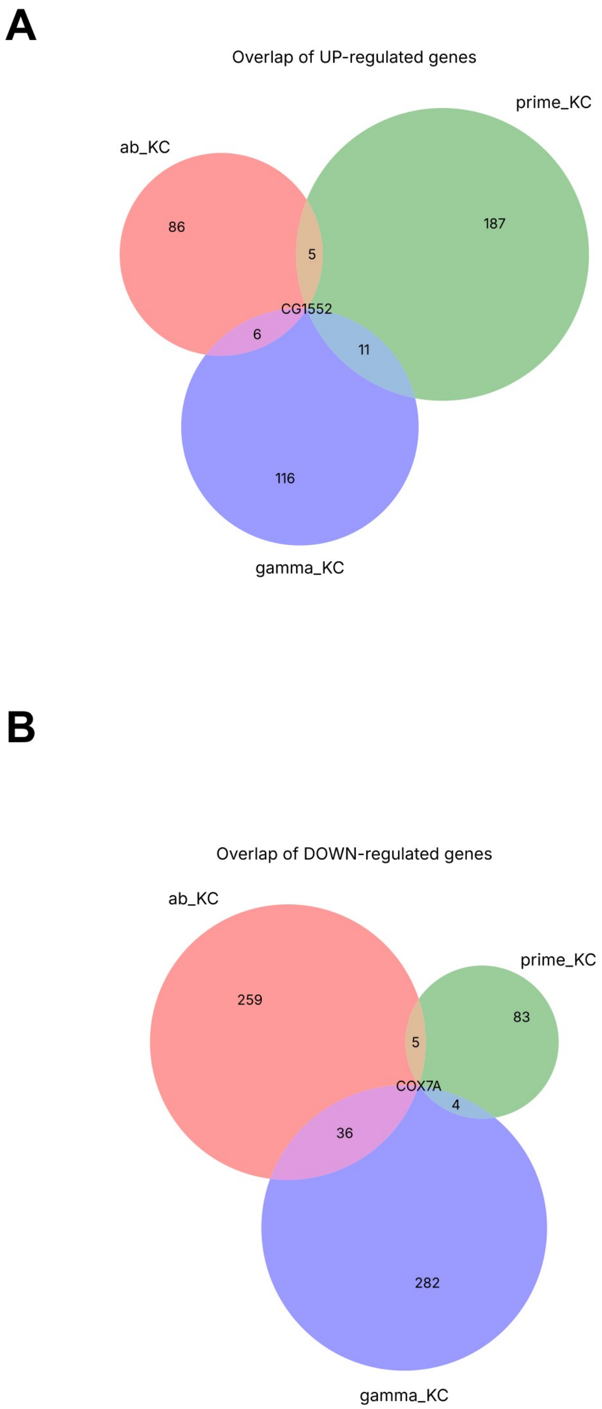


**Supplementary Figure 8. Overlap of DEGs between Kenyon cell types.** (**A**) Venn diagram showing the number of overlapping DEGs that are up-regulated in the Kenyon cell types of flies fed high [$Mg^{2+}$] diet compared with low [$Mg^{2+}$] diet. (**B**) Venn diagram showing the number of overlapping DEGs that are down-regulated in the Kenyon cell types of flies fed high [$Mg^{2+}$] diet.

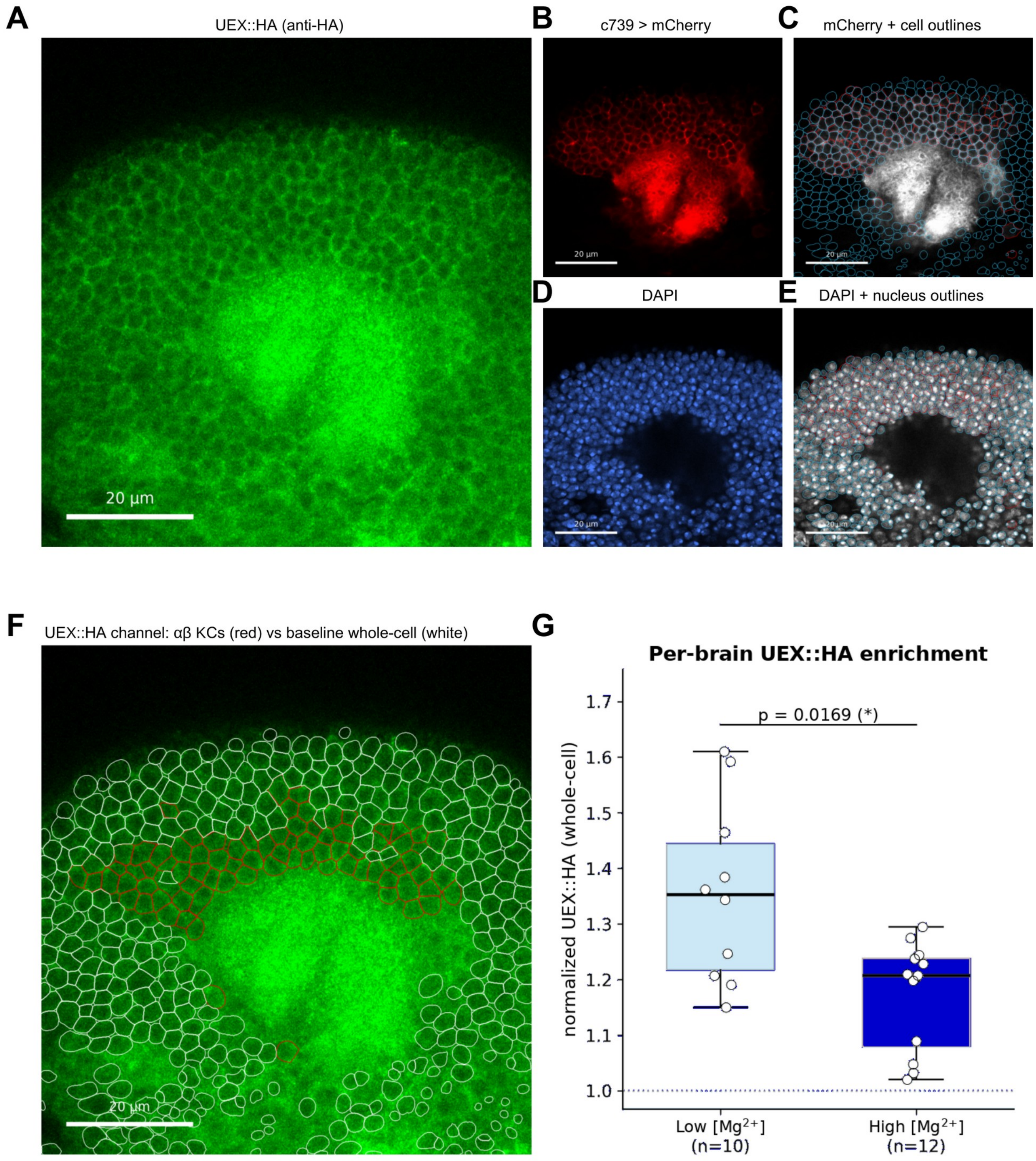


**Supplementary Figure 9. Quantification of Mg$^{2+}$ diet-dependent UEX::HA expression in αβ Kenyon cells.** Three-channel confocal z-stacks were acquired through the mushroom body Kenyon cell (KC) soma layer, comprising c739-GAL4-driven mCherry marked αβ KCs (red), anti-HA immunofluorescence detecting UEX::HA (green), and DAPI labeling nuclei (blue). Cell bodies and nuclei were segmented using Cellpose-SAM, and UEX::HA fluorescence was quantified independently in each optical section. (A to F) Representative optical section from a single brain illustrating the analysis workflow. (**A**) Anti-HA channel. (**B**) mCherry channel. (**C**) Cell-body segmentation overlaid on the mCherry channel; mCherry-positive αβ KCs are outlined in red and mCherry-negative cells in cyan. (**D**) DAPI channel. (**E**) Nuclear segmentation overlaid on the DAPI channel. (**F**) Anti-HA channel overlaid with the whole-cell compartments used for fluorescence quantification; mCherry-positive αβ KCs (signal population) are outlined in red and mCherry-negative cells (baseline population) in white. For each cell, the mean anti-HA fluorescence intensity was measured over the cell body. For baseline cells, the whole-cell compartment was estimated by expanding each DAPI-labeled nucleus with a perinuclear ring whose width equaled the difference between the median cell-body radius and the median nuclear radius of that optical section, excluding mCherry-positive pixels. Normalized UEX::HA enrichment was calculated for each optical section as the ratio of the mean anti-HA intensity in mCherry-positive αβ KCs to that in mCherry-negative baseline cells. Each brain was represented by the median normalized UEX::HA value across all analyzable optical sections. (**G**) Per-brain normalized UEX::HA enrichment in flies fed low (1 mM, $n$ = 10 brains) or high (80 mM, $n$ = 12 brains) [$Mg^{2+}$] diet. Each point represents one brain; boxes indicate the interquartile range with the median shown as the center line, whiskers indicate the full range, and the dotted line denotes a normalized value of 1 (no enrichment). UEX::HA enrichment in αβ KCs was greater in flies fed low [$Mg^{2+}$] diet than in those fed high [$Mg^{2+}$] diet (two-sided Mann-Whitney $U$ test, $P$ = 0.0169). Scale bars, 20 μm.

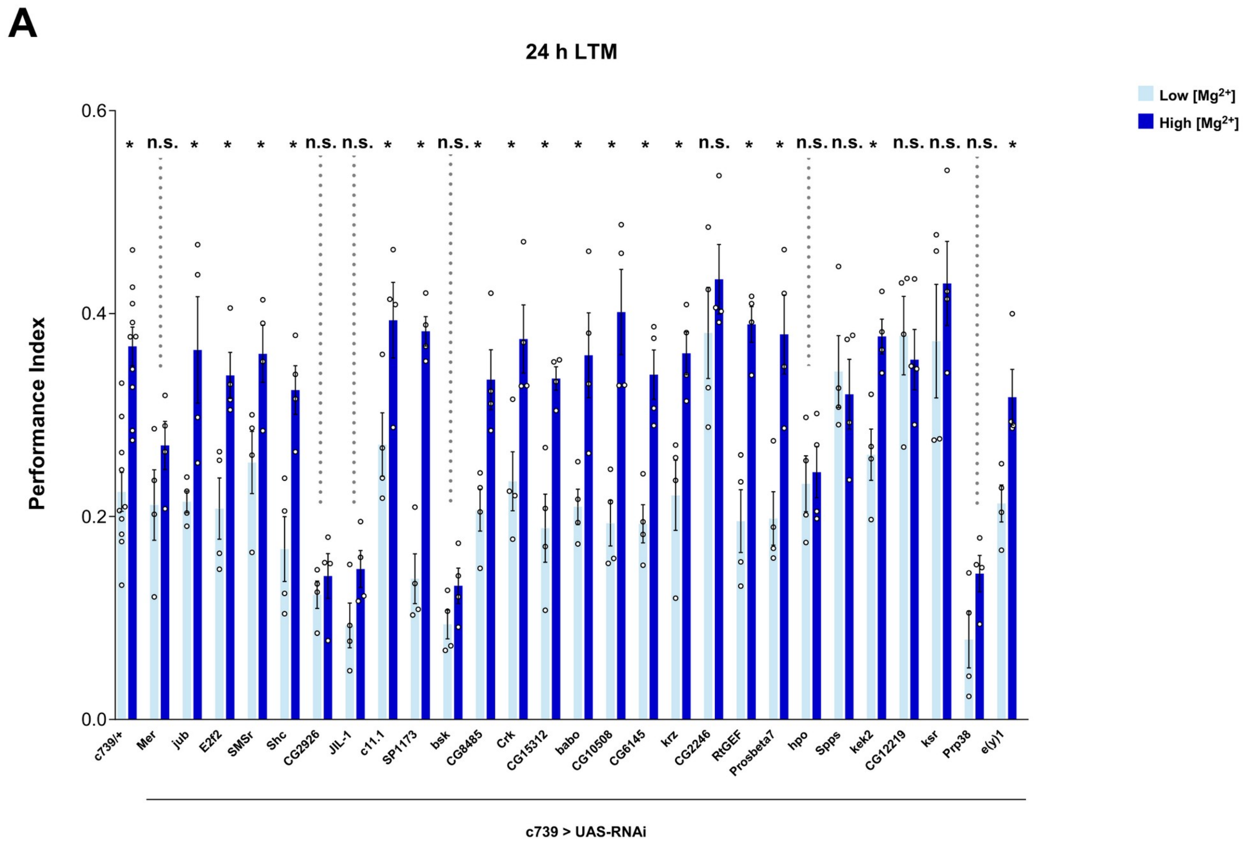


**Supplementary Figure 10. Screen for loss of $Mg^{2+}$-dependent memory enhancement.** Flies expressing RNAi knockdown of candidate DEGs under c739-GAL4 control were tested for appetitive 24 h memory after being fed high or low [$Mg^{2+}$] diet for 4 days. Knockdown of *Mer*, *CG2926*, *JIL-1*, *bsk*, *CG2246*, *hpo*, *Spps*, *CG12219*, *ksr* and *Prp38* produced loss of memory enhancement. Knockdown of the remaining candidate genes did not affect enhancement, with these flies retaining a significant increase in LTM when fed high [$Mg^{2+}$] diet (*$P < 0.05$, *t* test, $n = 4\text{-}8$).

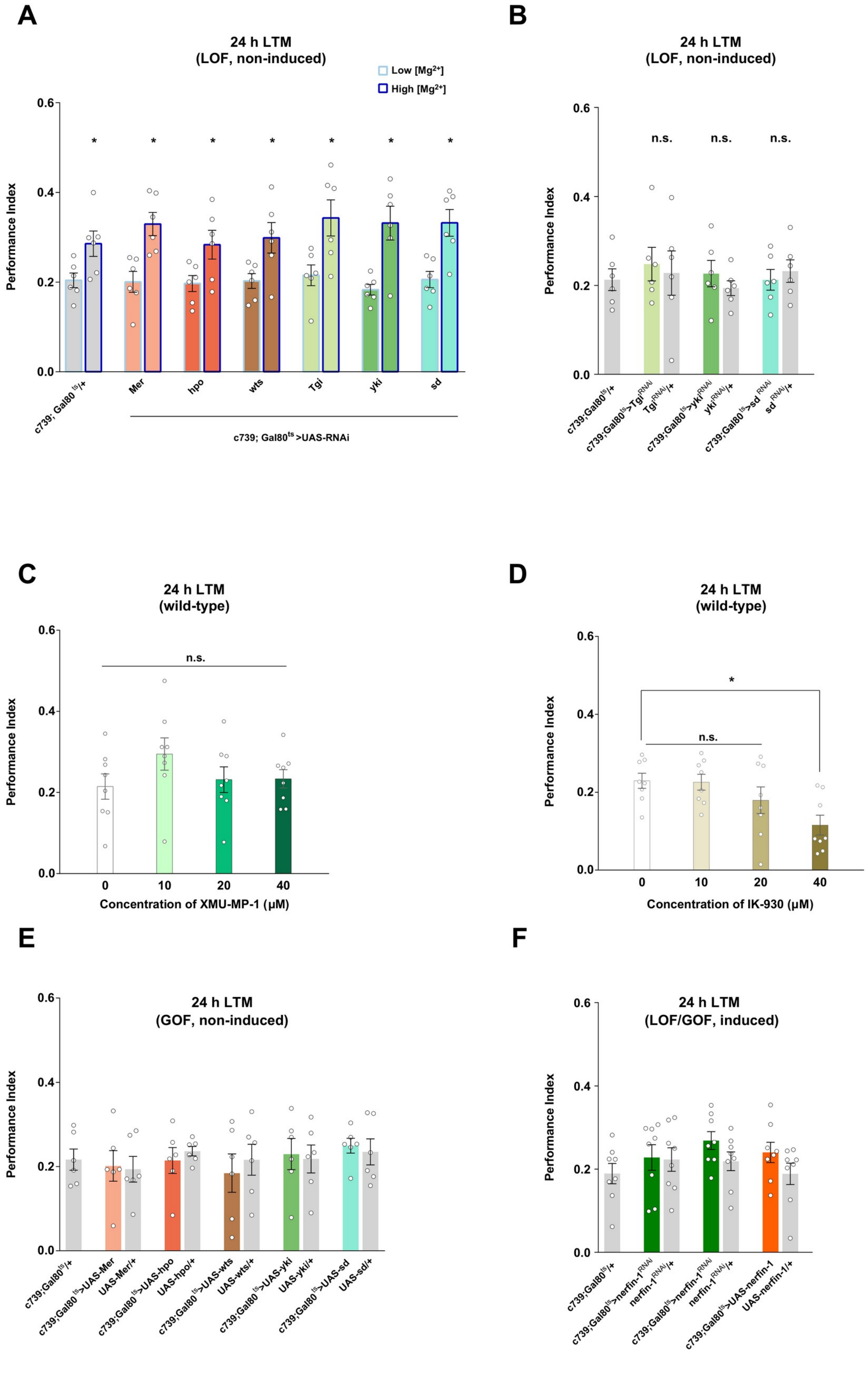
A
24 h LTM
(LOF, non-induced)
Low [Mg2+]
High [Mg2+]
Performance Index
0.0
0.2
0.4
0.6
*
c739; Gal80ts/+
Mer
hpo
wts
Tgi
yki
sd
c739; Gal80ts >UAS-RNAi
B
24 h LTM
(LOF, non-induced)
n.s.
c739;Gal80ts/+
c739;Gal80ts>TgiRNAi
TgiRNAi/+
c739;Gal80ts>ykiRNAi
ykiRNAi/+
c739;Gal80ts>sdRNAi
sdRNAi/+
C
24 h LTM
(wild-type)
0
10
20
40
Concentration of XMU-MP-1 (μM)
D
24 h LTM
(wild-type)
Concentration of IK-930 (μM)
E
24 h LTM
(GOF, non-induced)
c739;Gal80ts/+
c739;Gal80ts>UAS-Mer
UAS-Mer/+
c739;Gal80ts>UAS-hpo
UAS-hpo/+
c739;Gal80ts>UAS-wts
UAS-wts/+
c739;Gal80ts>UAS-yki
UAS-yki/+
c739;Gal80ts>UAS-sd
UAS-sd/+
F
24 h LTM
(LOF/GOF, induced)
c739;Gal80ts/+
c739;Gal80ts>nerfin-1RNAi
nerfin-1RNAi/+
c739;Gal80ts>nerfin-1RNAi
nerfin-1RNAi/+
c739;Gal80ts>UAS-nerfin-1
UAS-nerfin-1/+

**Supplementary Figure 11. Controls and pharmacological tests for involvement of Hippo pathway genes in LTM.** (**A**) Memory enhancement produced by high [$Mg^{2+}$] diet was not affected if RNAi knockdown of *Mer*, *hpo*, *wts*, *Tgi*, *yki* or *sd* was not induced (*$P < 0.05$, *t* test, $n = 6$). (**B**) Baseline LTM remained intact if RNAi knockdown of *Tgi*, *yki* or *sd* was not induced (ANOVA, $n = 6$). (**C**) Feeding flies 10, 20 or 40 μM XMU-MP-1 for 4 days did not affect LTM (ANOVA, $n = 8$). (**D**) Feeding flies 40 μM IK-930 for 4 days significantly disrupted LTM, whereas 10 μM or 20 μM IK-930 left LTM intact (*$P < 0.05$, ANOVA, $n = 8$). (**E**) LTM was not affected if overexpression of *Mer*, *hpo*, *wts*, *yki* or *sd* was not induced (ANOVA, $n = 6$). (**F**) Adult αβ KC knockdown of *nerfin-1* (two independent RNAi lines) or overexpression of *nerfin-1* neither enhanced or impaired LTM (ANOVA, $n = 8$).

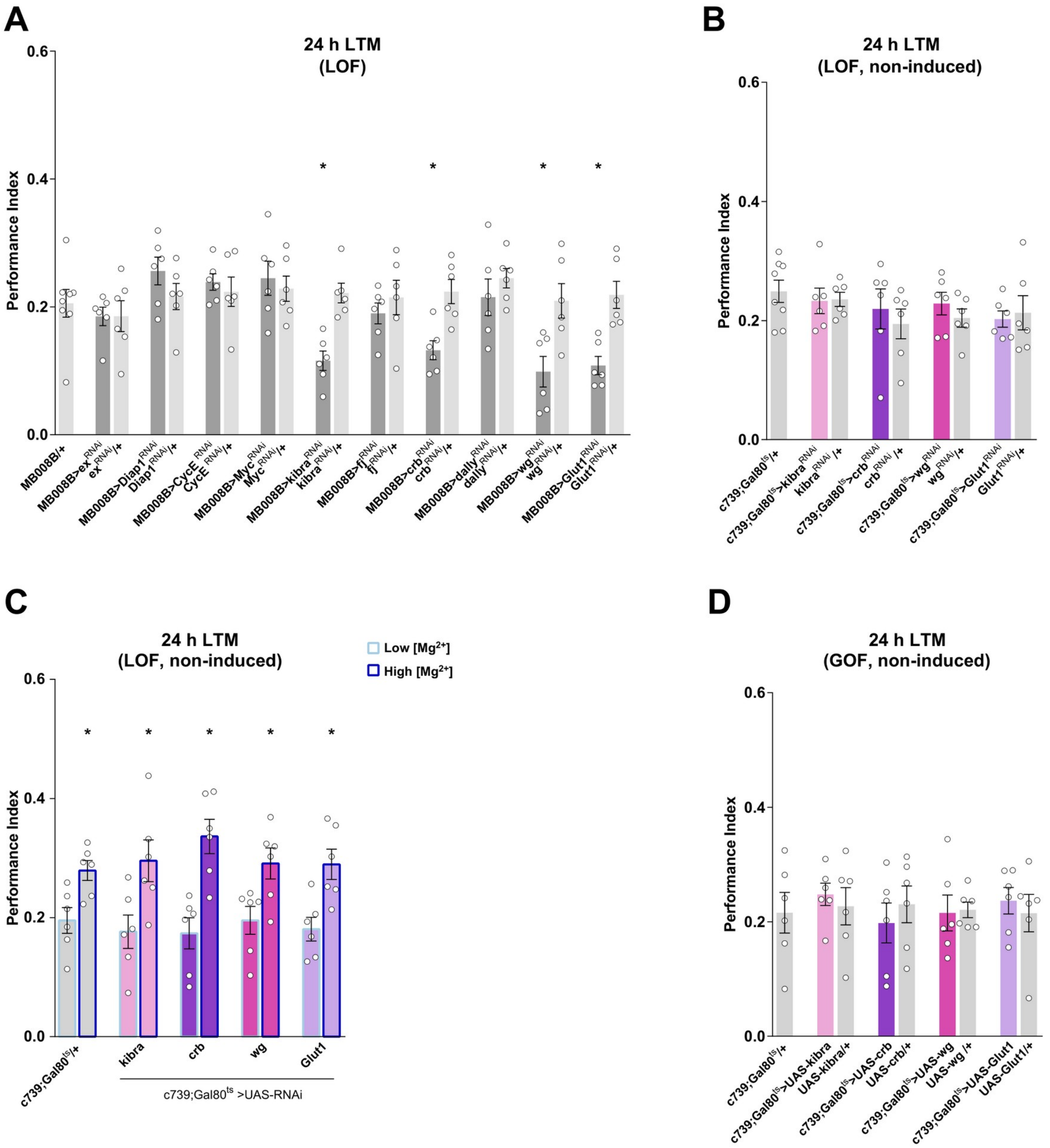


**Supplementary Figure 12. Controls for memory tests of Hippo pathway target genes.** (**A**) Baseline LTM was significantly impaired in flies with αβ KC knockdown of *kibra*, *crb*, *wg* or *Glut1*, but not of *ex*, *Diap1*, *CycE*, *Myc*, *fj* or *dally* (*$P < 0.05$, ANOVA, $n = 6$). (**B**) Baseline LTM remained intact if RNAi knockdown of *kibra*, *crb*, *wg* or *Glut1* was not induced (ANOVA, $n = 6$). (**C**) Memory enhancement produced by high [$Mg^{2+}$] diet was not affected if RNAi knockdown of *kibra*, *crb*, *wg* or *Glut1* was not induced (*$P < 0.05$, *t* test, $n = 6$). (**D**) LTM was not affected if overexpression of *kibra*, *crb*, *wg* or *Glut1* was not induced (ANOVA, $n = 6$).

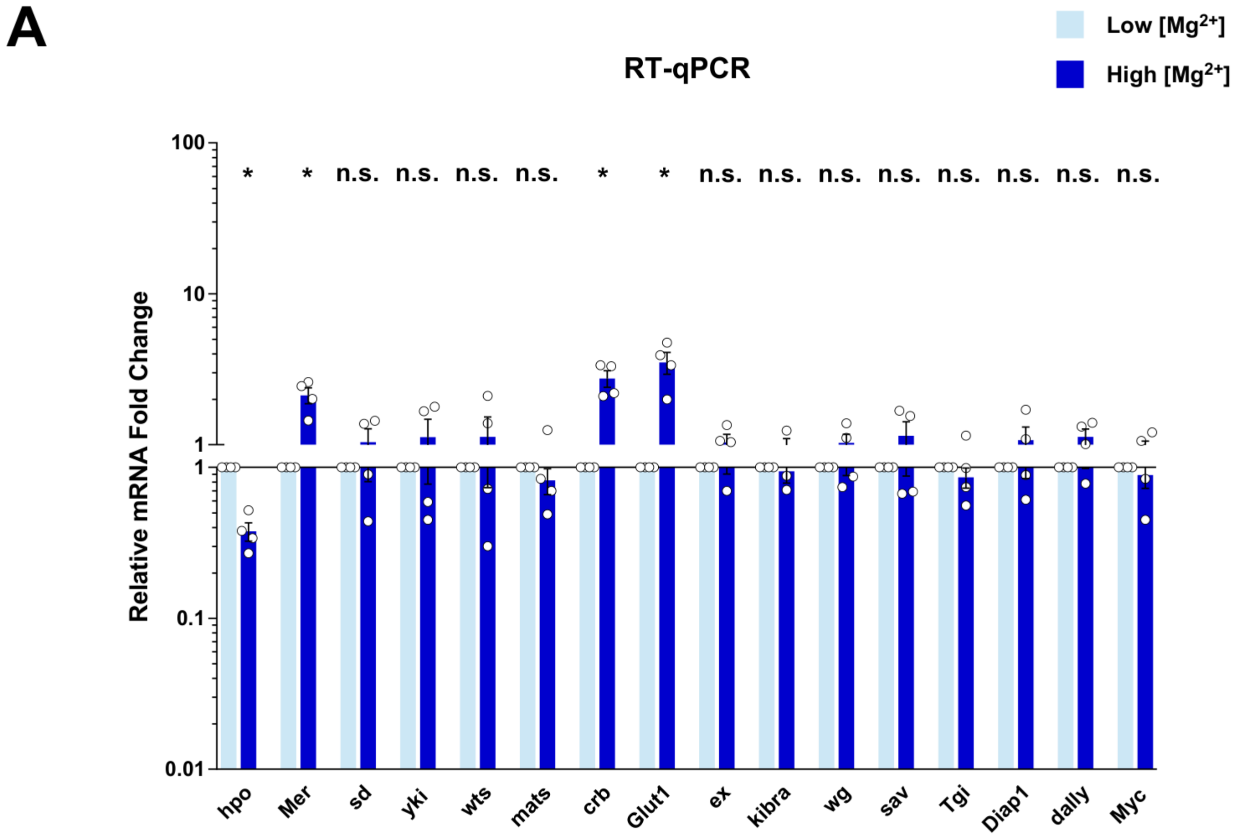


**Supplementary Figure 13. RT-qPCR measurement of Hippo pathway and target gene expression in αβ Kenyon cells.** (**A**) Relative mRNA fold changes of Hippo pathway and target genes in αβ KCs of flies fed high [$Mg^{2+}$] diet compared with those fed low [$Mg^{2+}$] diet. Each data point represents one biological replicate with 3 technical replicates. *$P < 0.05$, $t$ test, $n = 4$.

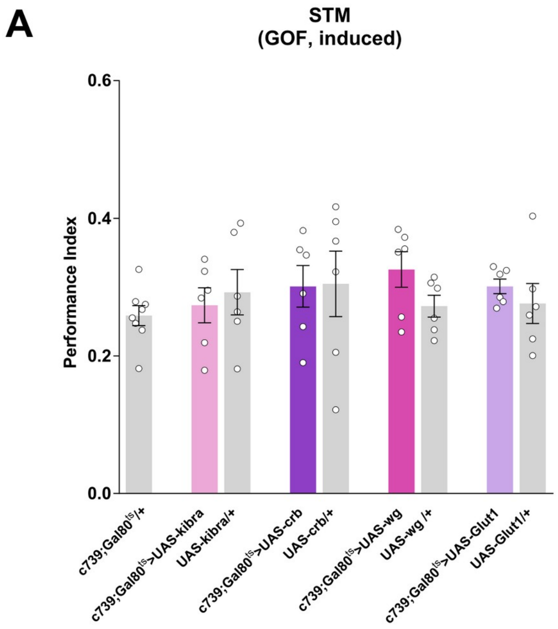


**Supplementary Figure 14. Short-term memory of flies with Hippo pathway target genes manipulated in adult αβ Kenyon cells.** (A) STM performance was unaffected by adult αβ Kenyon cell overexpression of *kibra*, *crb*, *wg* or *Glut1* (ANOVA, $n$ = 6-8).

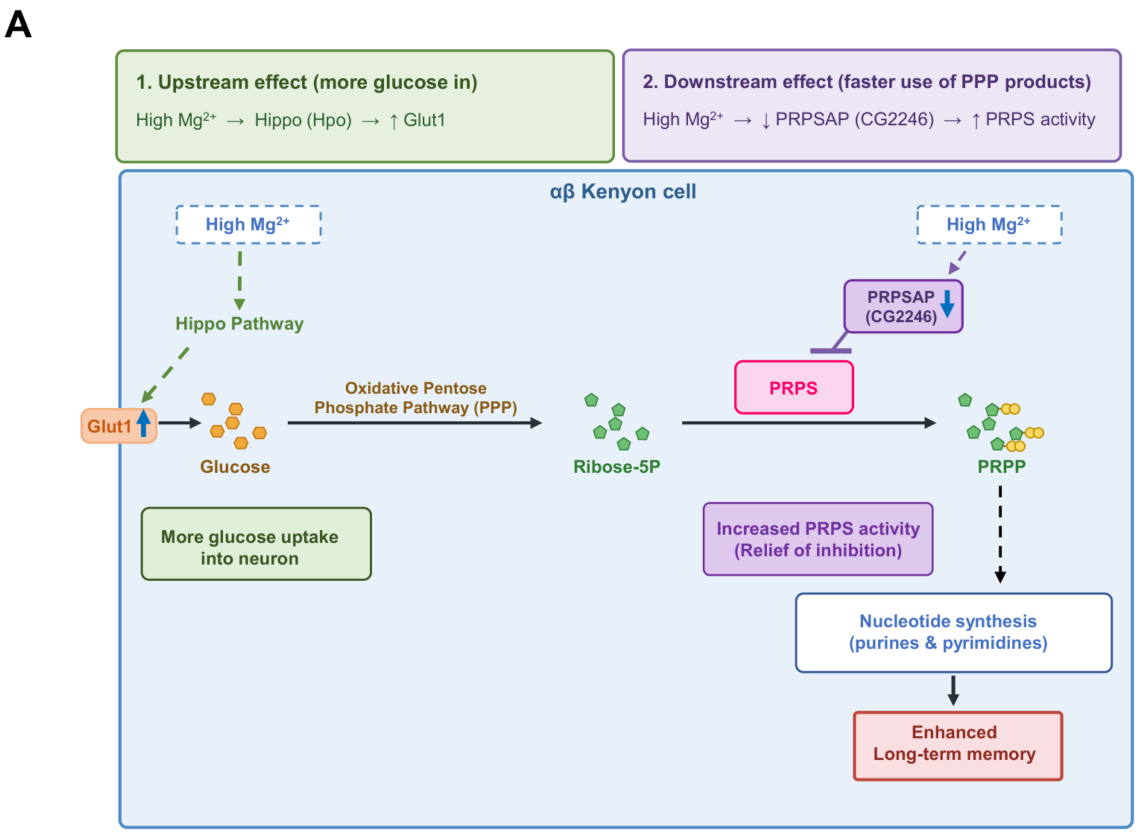


**Supplementary Figure 15. Proposed metabolic model for dietary $Mg^{2+}$-enhanced long-term memory in αβ Kenyon cells.** (A) Two parallel effects of high [$Mg^{2+}$] diet are proposed. (1) Upstream: high [$Mg^{2+}$] diet acts through the Hippo pathway (Hpo) to up-regulate *Glut1*, increasing glucose uptake into αβ KCs; glucose is metabolized by the oxidative pentose phosphate pathway (PPP) to ribose-5-phosphate (Ribose-5P). (2) Downstream: high [$Mg^{2+}$] diet down-regulates PRPSAP (*CG2246*), relieving its inhibition of phosphoribosyl pyrophosphate synthetase (PRPS) and increasing conversion of ribose-5-phosphate into phosphoribosyl pyrophosphate (PRPP), a precursor for purine and pyrimidine nucleotide synthesis that is proposed to support enhanced long-term memory.

**Supplementary Table 1. Key resources.** Antibodies, chemicals, commercial kits, deposited data, experimental models, oligonucleotides and software used in this study, with their sources and identifiers.

| REAGENT or RESOURCE | SOURCE | IDENTIFIER |
|---|---|---|
| Antibodies | | |
| Rat anti-HA | Merck | 11867423001; RRID:AB_390918 |
| Alexa Fluor 488-conjugated goat anti-rat | Invitrogen | A11006; RRID:AB_2534074 |
| Chemicals, Peptides, and Recombinant Proteins | | |
| UltraPure $H_2O$ | Life Technology | 10977-015 |
| NaCl | Merck | S6546-1L |
| $MgCl_2$ | Sigma | M1028 |
| XMU MP 1 | Fisher Scientific | 18787108 |
| IK-930 | Stratech | E1483-SEL |
| Sucrose | Sigma | S0389 |
| Mineral oil | Sigma | M5904 |
| 3-Octanol | Sigma-Aldrich | 218405 |
| 4-Methyl-Cyclohexanol | Sigma-Aldrich | 66360 |
| Paraformaldehyde | Fisher scientific | 15713 |
| Phosphate buffered saline tablets | Fisher scientific | 1282-1680 |
| Tris-HCl pH 7.4 | Merck | T2194-100ML |
| Tween-20 | Merck | P1379-25ML |
| Triton X-100 | Sigma | T9284 |
| NP-40 Alternative | Merck | 492016-100ML |
| 5% Digitonin | ThermoFisher | BN2006 |
| 2% Digitonin | promega | G9441 |
| DPBS (calcium and magnesium free) | Gibco | 14190–086 |
| HEPES solution | Merck | H0887-20ML |
| Schneider's medium | Gibco | 21720–001 |
| Papain | Sigma | P4762 |
| Collagenase | Sigma | C2674 |
| Pierce Protease inhibitor | ThermoFisher | A32953 |
| d(-)-2-amino-5-phosphonovaleric acid | Sigma | A8054 |
| DAPI Staining Solution | Abcam | ab228549 |
| 6,7-dinitroquinoxaline-2,3-dione | Sigma | D0540 |
| tetrodotoxin | Abcam | ab120055 |
| bovine serum albumin | New England Biolabs | B9000S |
| Critical Commercial Assay | | |
| Chromium Next GEM Single Cell 3' Kit v3.1, 4 rxns | 10x Genomics | PN-1000269 |
| SuperScript IV CellsDirect cDNA Synthesis Kit | Invitrogen | 11750150 |
| QIAquick PCR Purification Kit | Qiagen | 28106 |
| KAPA HiFi HotStart ReadyMix | Kapa Biosystems | KK2611 |
| Q5 High Fidelity Master Mix (2x) | NEB | M0492S |
| MinElute PCR Purification Kit (50) | Qiagen | 28004 |
| LigherCycler 480 SYBR Green I Master | Roche | 04707516001 |
| Experimental Models: Organisms/Strains | | |

| REAGENT or RESOURCE | SOURCE | IDENTIFIER |
|---|---|---|
| D. melanogaster, Canton-S | Lab stock (Dudai, Jan et al. 1976) | N/A |
| D. melanogaster, UAS-mCherry | Lab stock | N/A |
| D. melanogaster, c739-GAL4 | Lab stock (McGuire, Le et al. 2001) | N/A |
| D. melanogaster, tubP-GAL80$^{ts}$ | Lab stock (McGuire, Le et al. 2003) | N/A |
| D. melanogaster, MB008B | Bloomington Drosophila Stock Center | 68291; RRID:BDSC_68291 |
| D. melanogaster, UAS-*Mer*$^{RNAi}$ | Bloomington Drosophila Stock Center | 34958; RRID:BDSC_34958 |
| D. melanogaster, UAS-*jub*$^{RNAi}$ | Bloomington Drosophila Stock Center | 41938; RRID:BDSC_41938 |
| D. melanogaster, UAS-*E2f2*$^{RNAi}$ | Bloomington Drosophila Stock Center | 27995; RRID:BDSC_27995 |
| D. melanogaster, UAS-*SMSr*$^{RNAi}$ | Vienna Drosophila Resource Center | 39483; RRID:SCR_013805 |
| D. melanogaster, UAS-*Shc*$^{RNAi}$ | Bloomington Drosophila Stock Center | 66961; RRID:BDSC_66961 |
| D. melanogaster, UAS-*CG2926*$^{RNAi}$ | Bloomington Drosophila Stock Center | 34941; RRID:BDSC_34941 |
| D. melanogaster, UAS-*JIL-1*$^{RNAi}$ | Bloomington Drosophila Stock Center | 57293; RRID:BDSC_57293 |
| D. melanogaster, UAS-*c11.1*$^{RNAi}$ | Bloomington Drosophila Stock Center | 61204; RRID:BDSC_61204 |
| D. melanogaster, UAS-*SP1173*$^{RNAi}$ | Vienna Drosophila Resource Center | 109027; RRID:SCR_013805 |
| D. melanogaster, UAS-*bsk*$^{RNAi}$ | Bloomington Drosophila Stock Center | 53310; RRID:BDSC_53310 |
| D. melanogaster, UAS-*CG8485*$^{RNAi}$ | Bloomington Drosophila Stock Center | 60083; RRID:BDSC_60083 |
| D. melanogaster, UAS-*Crk*$^{RNAi}$ | Bloomington Drosophila Stock Center | 55277; RRID:BDSC_55277 |
| D. melanogaster, UAS-*CG15312*$^{RNAi}$ | Vienna Drosophila Resource Center | 101286; RRID:SCR_013805 |
| D. melanogaster, UAS-*babo*$^{RNAi}$ | Bloomington Drosophila Stock Center | 25933; RRID:BDSC_25933 |
| D. melanogaster, UAS-*CG10508*$^{RNAi}$ | Vienna Drosophila Resource Center | 102744; RRID:SCR_013805 |
| D. melanogaster, UAS-*CG6145*$^{RNAi}$ | Bloomington Drosophila Stock Center | 57753; RRID:BDSC_57753 |
| D. melanogaster, UAS-*krz*$^{RNAi}$ | Bloomington Drosophila Stock Center | 33880; RRID:BDSC_33880 |
| D. melanogaster, UAS-*CG2246*$^{RNAi}$ | Vienna Drosophila Resource Center | 110389; RRID:SCR_013805 |
| D. melanogaster, UAS-*RtGEF*$^{RNAi}$ | Vienna Drosophila Resource Center | 105093; RRID:SCR_013805 |
| D. melanogaster, UAS-*Prosbeta7*$^{RNAi}$ | Bloomington Drosophila Stock Center | 34812; RRID:BDSC_34812 |
| D. melanogaster, UAS-*hpo*$^{RNAi}$ | Bloomington Drosophila Stock Center | 33614; RRID:BDSC_33614 |
| D. melanogaster, UAS-*Spps*$^{RNAi}$ | Bloomington Drosophila Stock Center | 62185; RRID:BDSC_62185 |

| REAGENT or RESOURCE | SOURCE | IDENTIFIER |
|---|---|---|
| D. melanogaster, UAS-*kek2*$^{RNAi}$ | Bloomington Drosophila Stock Center | 31874; RRID:BDSC_31874 |
| D. melanogaster, UAS-*CG12219*$^{RNAi}$ | Bloomington Drosophila Stock Center | 28000; RRID:BDSC_28000 |
| D. melanogaster, UAS-*ksr*$^{RNAi}$ | Bloomington Drosophila Stock Center | 32937; RRID:BDSC_32937 |
| D. melanogaster, UAS-*Prp38*$^{RNAi}$ | Vienna Drosophila Resource Center | 110282; RRID:SCR_013805 |
| D. melanogaster, UAS-*e(y)1*$^{RNAi}$ | Bloomington Drosophila Stock Center | 32345; RRID:BDSC_32345 |
| D. melanogaster, UAS-*ex*$^{RNAi}$ | Bloomington Drosophila Stock Center | 28703; RRID:BDSC_28703 |
| D. melanogaster, UAS-*sav*$^{RNAi}$ | Bloomington Drosophila Stock Center | 32965; RRID:BDSC_32965 |
| D. melanogaster, UAS-*wts*$^{RNAi}$ | Bloomington Drosophila Stock Center | 27662; RRID:BDSC_27662 |
| D. melanogaster, UAS-*mats*$^{RNAi}$ | Bloomington Drosophila Stock Center | 34959; RRID:BDSC_34959 |
| D. melanogaster, UAS-*Tgi*$^{RNAi}$ | Vienna Drosophila Resource Center | 31340; RRID:SCR_013805 |
| D. melanogaster, UAS-*yki*$^{RNAi}$ | Bloomington Drosophila Stock Center | 34067; RRID:BDSC_34067 |
| D. melanogaster, UAS-*sd*$^{RNAi}$ | Vienna Drosophila Resource Center | 101497; RRID:SCR_013805 |
| D. melanogaster, UAS-*Diap1*$^{RNAi}$ | Bloomington Drosophila Stock Center | 33975; RRID:BDSC_33975 |
| D. melanogaster, UAS-*CycE*$^{RNAi}$ | Bloomington Drosophila Stock Center | 38902; RRID:BDSC_38902 |
| D. melanogaster, UAS-*Myc*$^{RNAi}$ | Bloomington Drosophila Stock Center | 43962; RRID:BDSC_43962 |
| D. melanogaster, UAS-*kibra*$^{RNAi}$ | Bloomington Drosophila Stock Center | 28683; RRID:BDSC_28683 |
| D. melanogaster, UAS-*fj*$^{RNAi}$ | Bloomington Drosophila Stock Center | 34323; RRID:BDSC_34323 |
| D. melanogaster, UAS-*crb*$^{RNAi}$ | Bloomington Drosophila Stock Center | 27697; RRID:BDSC_27697 |
| D. melanogaster, UAS-*dally*$^{RNAi}$ | Bloomington Drosophila Stock Center | 33952; RRID:BDSC_33952 |
| D. melanogaster, UAS-*wg*$^{RNAi}$ | Bloomington Drosophila Stock Center | 32994; RRID:BDSC_32994 |
| D. melanogaster, UAS-*Glut1*$^{RNAi}$ | Vienna Drosophila Resource Center | 101365; RRID:SCR_013805 |
| D. melanogaster, UAS-*nerfin-1*$^{RNAi}$ | Bloomington Drosophila Stock Center | 28324; RRID:BDSC_28324 |
| D. melanogaster, UAS-*nerfin-1*$^{RNAi}$ | Vienna Drosophila Resource Center | 101631; RRID:SCR_013805 |
| D. melanogaster, UAS-*Glut1* | Lab stock (Niccoli, Cabecinha et al. 2016) | N/A |
| D. melanogaster, UAS-*Mer* | Bloomington Drosophila Stock Center | 600495; RRID:BDSC_600495 |
| D. melanogaster, UAS-*hpo* | Bloomington Drosophila Stock Center | 600509; RRID:BDSC_600509 |
| D. melanogaster, UAS-*wts* | Bloomington Drosophila Stock Center | 600510; RRID:BDSC_600510 |

| REAGENT or RESOURCE | SOURCE | IDENTIFIER |
|---|---|---|
| D. melanogaster, UAS-*yki* | Bloomington Drosophila Stock Center | 600526; RRID:BDSC_600526 |
| D. melanogaster, UAS-*sd* | Bloomington Drosophila Stock Center | 9374; RRID:BDSC_9374 |
| D. melanogaster, UAS-*kibra* | Bloomington Drosophila Stock Center | 600581; RRID:BDSC_600581 |
| D. melanogaster, UAS-*crb* | Bloomington Drosophila Stock Center | 5544; RRID:BDSC_5544 |
| D. melanogaster, UAS-*wg* | Bloomington Drosophila Stock Center | 5918; RRID:BDSC_5918 |
| D. melanogaster, UAS-*nerfin-1* | Bloomington Drosophila Stock Center | 67385; RRID:BDSC_67385 |
| D. melanogaster, TRiP-attP2 | Bloomington Drosophila Stock Center | 36303; RRID:BDSC_36303 |
| D. melanogaster, TRiP-attP40 | Bloomington Drosophila Stock Center | 36304; RRID:BDSC_36304 |
| D. melanogaster, *uex::HA* | Lab stock (Wu, Funato et al. 2020) | N/A |
| Oligonucleotides | | |
| TGTACGACACGGTCAAGGG | This study | qPCR of SdhA Forward |
| TTCTCCAGCTCAATGACAGCC | This study | qPCR of SdhA Reverse |
| TTTACTCGCAGGACCTCCAC | This study | qPCR of *Mer* Forward |
| GCGGCGCATATATAGATCGT | This study | qPCR of *Mer* Reverse |
| CGCCGGTATATGAGAACTTGG | This study | qPCR of *jub* Forward |
| CCGGGACTTTGAGGCTGAG | This study | qPCR of *jub* Reverse |
| GCCAATCAAGTCTCCGTCGAA | This study | qPCR of *E2f2* Forward |
| AACTTCTGTGTGAGCAGGACC | This study | qPCR of *E2f2* Reverse |
| AATTGCCGCCAAATTCAGTCG | This study | qPCR of *SMSr* Forward |
| CGCACGGGTCAGTTCTATTAAA | This study | qPCR of *SMSr* Reverse |
| TTAGCGCGAGAGTGCATCAAT | This study | qPCR of *Shc* Forward |
| TGAAGTTGGTTAATCTCCGCTTG | This study | qPCR of shc Reverse |
| AGTTCCGACATCAGCAGTAACG | This study | qPCR of *CG2926* Forward |
| AGGCCGCGCAGAATATGTG | This study | qPCR of *CG2926* Reverse |
| ACGGTGGTCCAGAAGCGAAA | This study | qPCR of *JIL-1* Forward |
| CCTCCAGTACCACTCTCTC | This study | qPCR of *JIL-1* Reverse |
| CCGAAGATGACCGATCAAACA | This study | qPCR of *c11.1* Forward |

| REAGENT or RESOURCE | SOURCE | IDENTIFIER |
|---|---|---|
| ATGCTCGGGATTCTTGGTGAG | This study | qPCR of *c11.1* Reverse |
| ATGGGTTTCATTGCGGAGC | This study | qPCR of *SP1173* Forward |
| TGCAAAACATGCAAAGCTGTAAG | This study | qPCR of *SP1173* Reverse |
| GCTCAGCACCAACACTACAC | This study | qPCR of *bsk* Forward |
| GCGCATACTATTCCTTGGGCA | This study | qPCR of *bsk* Reverse |
| CAAGGTGGCGGTCAAGGTAG | This study | qPCR of *CG8485* Forward |
| ATCGCACTTCCTGGAACAGAT | This study | qPCR of *CG8485* Reverse |
| TTCGTTGGCAGCGATCAAGAT | This study | qPCR of *Crk* Forward |
| TACGCGCAGTCCACCATTG | This study | qPCR of *Crk* Reverse |
| CCAAGTGCTCACGGTACTCAG | This study | qPCR of *CG15312* Forward |
| ATGCACGGTATGCTCACATTG | This study | qPCR of *CG15312* Reverse |
| CTACCAGATTATGTGCCACAC | This study | qPCR of *babo* Forward |
| TACTGGTGCCCGTGAAGCAA | This study | qPCR of *babo* Reverse |
| GCAAATGCGGATAACAACGTG | This study | qPCR of *CG10508* Forward |
| TGTGGCTGTTGCATATCTCTTTT | This study | qPCR of *CG10508* Reverse |
| AAGTGCATCAACAGTACGCC | This study | qPCR of CG6145 Forward |
| GCAGACGATAAAGTCAATCCGAT | This study | qPCR of CG6145 Reverse |
| CTCCAGCACCAATGTAACAGC | This study | qPCR of *krz* Forward |
| GCTCTTCTTAAACACCCGAGTC | This study | qPCR of *krz* Reverse |
| GATTCCGTTAGAGGCAAGGAC | This study | qPCR of *CG2246* Forward |
| AGGCCATAATCAGAAGCTCCAT | This study | qPCR of *CG2246* Reverse |
| AACGATAAGACTGGCTGGTTTC | This study | qPCR of *RtGEF* Forward |
| TATGGTCTCCGTTAGAGGCAG | This study | qPCR of *RtGEF* Reverse |
| ACCACAATGGGACCCTATGGA | This study | qPCR of *Prosbeta7* Forward |
| GACGCGCTCAATGTTTTGGTA | This study | qPCR of *Prosbeta7* Reverse |
| CCATTATGCAACAATGCGACTC | This study | qPCR of *hpo* Forward |
| GATCCACAGGTCATACTGCTTG | This study | qPCR of *hpo* Reverse |

| REAGENT or RESOURCE | SOURCE | IDENTIFIER |
|---|---|---|
| TTCCACCAAATGGAGCTAGTGC | This study | qPCR of *Spps* Forward |
| CTCCGCCTGCAAACTGAGG | This study | qPCR of *Spps* Reverse |
| TGGTCGAGCTGGATCTCAG | This study | qPCR of *kek2* Forward |
| AGGCGCATTAGAGAGCTGTAG | This study | qPCR of *kek2* Reverse |
| CGATTCACTTGTACCTCTGCC | This study | qPCR of *CG12219* Forward |
| GGAAGCTCTCCAGTTGCGAG | This study | qPCR of *CG12219* Reverse |
| CGAAGATGCGGCTAAATGAGC | This study | qPCR of *ksr* Forward |
| GCAGCCATTGCCTCAGTTC | This study | qPCR of *ksr* Reverse |
| ATTCAGCCGGAAAAGGACATC | This study | qPCR of *Prp38* Forward |
| AAGGCTCCCAATGCTCGAAC | This study | qPCR of *Prp38* Reverse |
| GGAGCTGAATGTCCAGGAGTA | This study | qPCR of *e(y)1* Forward |
| GCAGGTTACATAGCGGAAGGT | This study | qPCR of *e(y)1* Reverse |
| CACGCACAAATTCCGCTTACT | This study | qPCR of *sd* Forward |
| TTGAACGAATAGATGCCGGTG | This study | qPCR of *sd* Reverse |
| ATCAGCCCCATTCAGTTGAAC | This study | qPCR of *yki* Forward |
| CCTCCCACTGCGTAGATTTTGTA | This study | qPCR of *yki* Reverse |
| TTGTACTTCCTAGTCGATGCCA | This study | qPCR of *ex* Forward |
| CGCCATCTATGAGCACAGCC | This study | qPCR of *ex* Reverse |
| ATCAAGCAGGACCTAACCCG | This study | qPCR of *wts* Forward |
| GCCGTGTATCGCAGAGGTGT | This study | qPCR of *wts* Reverse |
| CCAAGAAGAACATCCCGGAAG | This study | qPCR of *mats* Forward |
| TGGTACTCGTATTTCGGACCAG | This study | qPCR of *mats* Reverse |
| GGATCTGCCCAAGGTGTTTA | This study | qPCR of *crb* Forward |
| AGTTCTTTCCAATGATGCCG | This study | qPCR of *crb* Reverse |
| CCAAGTCGAGGGCAAACAGAA | This study | qPCR of *wg* Forward |
| TGGATCGCTGGGTCCATGTA | This study | qPCR of *wg* Reverse |
| CAGAGATTGCTACACAAAGCCG | This study | qPCR of *kibra* Forward |

| REAGENT or RESOURCE | SOURCE | IDENTIFIER |
|---|---|---|
| TTCGGATCGTAGGATTCCTCC | This study | qPCR of *kibra* Reverse |
| GCTATGCCACACACGAATACG | This study | qPCR of *Glut1* Forward |
| GACTATGGCTGTGCTTACGC | This study | qPCR of *Glut1* Reverse |
| CTGTGCAACCGAAAACCGAC | This study | qPCR of *sav* Forward |
| GCTGATCGCTCCACACATTGA | This study | qPCR of *sav* Reverse |
| GACCCAGCGTCATTACCCAG | This study | qPCR of tgi Forward |
| CACATCGCCATATCTGTAGAGTG | This study | qPCR of tgi Reverse |
| ATACCACAGCCCTTTCAGCC | This study | qPCR of *fj* Forward |
| CCCCCAACCGCTTGAGTAAT | This study | qPCR of *fj* Reverse |
| CCCCAGTATCCCGAATACGC | This study | qPCR of *Diap1* Forward |
| TCTGTTTCAGGTTCCTCGGC | This study | qPCR of *Diap1* Reverse |
| AGCCTCCATCAGCTAAGCG | This study | qPCR of *CycE* Forward |
| GCAACCGATGACAGATTGCC | This study | qPCR of *CycE* Reverse |
| GCCGGAATGTTCGAGCAAC | This study | qPCR of *dally* Forward |
| ACGTGACTTTGGAACTGTTTGG | This study | qPCR of *dally* Reverse |
| ATGGCCCTTTACCGCTCTG | This study | qPCR of *Myc* Forward |
| CGAGATCACTCTGAATGGTGGA | This study | qPCR of *Myc* Reverse |
| Software and Algorithms | | |
| Prism 6.0 | GraphPad | https://www.graphpad.com; RRID:SCR_002798 |
| R v4.2 | R Development Core Team, 2008 | http://www.R-project.org/; RRID:SCR_001905 |
| Rstudio v2022.07 | RStudio, PBC | www.rstudio.com; RRID:SCR_000432 |
| Python v3.10 | The Python Software Foundation | https://www.python.org/; RRID:SCR_008394 |
| Jupyter-lab v0.30 | The Jupyter Book community | https://jupyter.org/; RRID:SCR_018315 |
| CellRanger v4.1 | 10X Genomics | https://support.10xgenomics.com/singlecell-gene-expression/software/overview/welcome; RRID:SCR_017344 |

| REAGENT or RESOURCE | SOURCE | IDENTIFIER |
| --- | --- | --- |
| SingularityCE v3.9 | Sylabs (Kurtzer, Sochat et al. 2017) | https://sylabs.io/ ; RRID:SCR_016248 |
| Seurat v4.0 | Satija Lab (Hao, Hao et al. 2021) | https://satijalab.org/seurat/articles/get_started.html; RRID:SCR_016341 |
| DoubletFinder v2.0 | Gartner Lab (McGinnis, Murrow et al. 2019) | https://github.com/chris-mcginnis-ucsf/DoubletFinder; RRID:SCR_018771 |
| Muscat | Robinson Lab (Crowell, Soneson et al. 2020) | https://github.com/HelenaLC/muscat-comparison |
| DAVID | DAVID Bioinformatics Team (Sherman, Hao et al. 2022) | https://david.ncifcrf.gov/ ; RRID:SCR_001881 |
| Cellpose | Stringer Lab (Stringer, Wang et al. 2021) | https://www.cellpose.org/ ; RRID:SCR_021105 |
| Other | | |
| 10 mm CellTrix strainer | Sysmex | 04-0042-2314 |
| Fuchs-Rosental hemocytometer | VWR | 631–1096 |

**Supplementary Data**

**Supplementary Data 1. (separate file)** DEGs identified by the Muscat differential expression analysis (DEGs_Muscat.xlsx). Tables of DEGs identified by the Muscat DE method for each annotated cell type.

**Supplementary Data 2. (separate file)** Integrated differential expression results (DEGs_Summary.xlsx). Tables of the merged DE results across the three methods for each annotated cell type.

**Supplementary Data 3. (separate file)** DEGs identified by the Cocktail differential expression analysis (DEGs_Cocktail.xlsx). Tables of DEGs identified by the Cocktail DE method for each annotated cell type.

**Supplementary Data 4. (separate file)** Kinase-related DEGs in αβ Kenyon cells (ab_KC_DEG_kinase-related.xlsx). Table of the 22 kinase-related DEGs identified in αβ Kenyon cells.

**Supplementary Data 5. (separate file)** Prioritized DEGs in αβ Kenyon cells (ab_KC_DEG_Essential.xlsx). Table of the 40 prominent DEGs in αβ Kenyon cells, combining the consensus and kinase-related groups. Genes selected for RT-qPCR validation are highlighted in bold.

**Supplementary Data 6. (separate file)** DEGs identified by the DirectE differential expression analysis (DEGs_DirectE.xlsx). Tables of DEGs identified by the DirectE DE method for each annotated cell type.

**Supplementary Data 7. (separate file)** Sensory acuity tests for Hippo pathway gene manipulations (sensory acuity_Hippo signalling.xlsx). Tables of odor avoidance and sugar preference test results.

**Supplementary Data 8. (separate file)** Sensory acuity tests for Hippo pathway target gene manipulations (sensory acuity_Hippo targets.xlsx). Tables of odor avoidance and sugar preference test results.
